# Opsin gene expression plasticity and spectral sensitivity in male damselflies could mediate female colour morph detection

**DOI:** 10.1101/2023.07.31.551331

**Authors:** Natalie S. Roberts, Erik I. Svensson, Marjorie A. Liénard

## Abstract

The visual systems of Odonata are characterized by many opsin genes, which form the primary light-sensitive photopigments of the eye. Female-limited colour polymorphisms are also common in Odonata, with one morph typically exhibiting male-like (androchrome) colouration and one or two morphs exhibiting female-specific colouration (gynochromes). These colour polymorphisms are thought to be maintained by frequency-dependent sexual conflict, in which males form search images for certain morphs, causing disproportionate mating harassment. Here, we investigate opsin sensitivity and gene expression plasticity in mate-searching males of the damselfly *Ischnura elegans* during adult maturation and across populations with different female morph frequencies. We find evidence for opsin-specific plasticity in relative and proportional opsin mRNA expression, suggesting changes in opsin regulation and visual sensitivity during sexual maturation. In particular, expression of the long-wavelength sensitive opsin LWF2 changed over development and varied between populations with different female morph frequencies. UV-Vis analyses indicate that short- and long-wavelength opsins absorb wavelength of light between 350 and 650 nm. Assuming opponency between photoreceptors with distinct short- and long-wavelength sensitivities, these sensitivities suggest male spectral visual discrimination ability of andromorph and gynomorph females. Overall, our results suggest that opsin sensitivity and expression changes contribute to visual tuning that could impact conspecific discrimination.

## INTRODUCTION

Opsins are G-protein coupled receptors that, when bound with a vitamin-A derived chromophore, form light sensitive visual pigments responsible for colour vision in animals (Terakita, 2005). Many insects possess three classes of opsin proteins: ultraviolet sensitive (UVS), short-wavelength sensitive (SWS), and long-wavelength sensitive (LWS) with maximum sensitivity to wavelengths (λ_max_) around 350 nm, 440 nm, and 530 nm, respectively (Briscoe & Chittka, 2001; van der Kooi et al., 2021). Photoreceptor spectral sensitivity depends mainly on the absorption spectrum of expressed opsins, which can vary due to opsin gains or losses, structural changes, or expression level differences (Castiglione et al., 2023; Liénard et al., 2021; Sharkey et al., 2023). Spectral sensitivity can also be modified by screening and filtering mechanisms, which are relatively widespread in insects (van der Kooi et al., 2021). The visual system is also shaped by environmental factors, behaviours, and associated selection pressures. For example, loss of functional opsin genes correlates with nocturnal lifestyles in mammals (Jacobs, 2013), while opsin duplications in other species contribute to the regaining of spectral sensitivity (Rossetto et al., 2023; Sharkey et al., 2023). In several insect species, functional changes amongst duplicated opsins likely improves detection and/or discrimination of environmental and sexual signals (Liénard et al., 2021; McCulloch et al., 2017).

Opsin gene expression is typically not fixed, but changes plastically over individual ontogeny or in response to environmental factors and lightning conditions (Chang et al., 2021; Fuller et al., 2010; Shand et al., 2008). For example, changes in opsin expression have been associated with light exposure in honey bees and ants (Sasagawa et al., 2003; Yilmaz et al., 2016), and in “choosy” relative to “non-choosy” females of the butterfly *Bicyclus anynana* (Everett et al., 2012).

Damselflies and dragonflies (Insecta: Odonata) are colourful diurnal flying insects that possess large conspicuous eyes comprising thousands of ommatidia (Fig. 1A). Odonates inhabit a wide array of visual environments, having both aquatic and terrestrial life stages, and they strongly rely on vision for a variety of behaviours including predator avoidance, prey capture, and mate searching (Bybee et al., 2012; Corbet, 1999; Labhart & Nilsson, 1995). These insects are therefore excellent model organisms to study evolutionary and functional aspects of colour vision. Like many other insects, odonates express UVS, SWS, and LWS opsins in their eyes. However, duplications of the SWS and LWS opsin classes have resulted in remarkable genetic diversity, with between 11 – 30 visual opsin genes identified across several families (Futahashi et al., 2015). Electrophysiological recordings of visual sensitivity in the eyes of some species indicate photoreceptor spectral curves ranging from UV to red light (Bybee et al., 2012). While it is not clear how their abundant opsin copies function in vision, visual stimuli are clearly important for odonate behaviour. For example, comparisons of visual and olfactory cues suggested that vision is primarily used in mate choice in several species of damselflies (Rebora et al., 2018; Winfrey & Fincke, 2017). Colouration is also an important signal indicating female sexual maturity (Svensson et al., 2020; Takahashi & Watanabe, 2011; Van Gossum et al., 2011; Willink et al., 2019). Males can discriminate between immature and mature females, and they direct more mating attempts to mature than immature females (Huang et al., 2014; Van Gossum et al., 2001a, 2011; Willink et al., 2019). Experimental manipulation of age-specific colouration in *Ischnura* damselflies has further confirmed that vision is a primary discrimination cue between immature and mature females (Willink et al., 2019).

**Figure 1.**
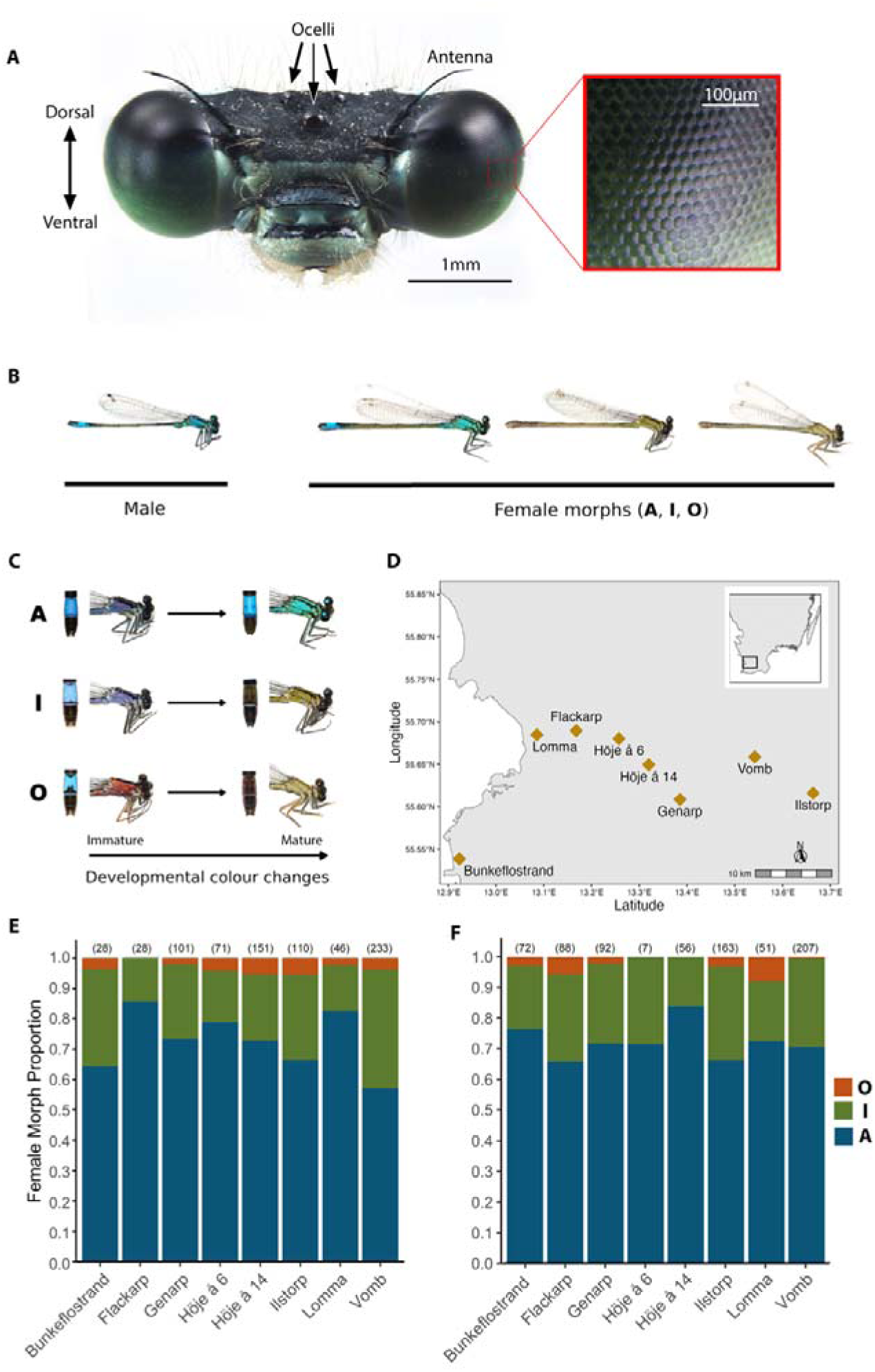
Head and eyes, colour polymorphism, and female morph frequencies from eight populations in southern Sweden for the damselfly *Ischnura elegans*. (A) Head of male *I. elegans*, showing the large compound eyes and three dorsally situated ocelli. Box shows magnified view of the compound eye depicting individual ommatidial units. (B) Mature male *I. elegans* and the three female-limited colour morphs in their sexual mature colour phases. (C) Females undergo ontogenetic colour changes in the distal abdomen segments (left photo) and thorax (right photo) between immature and sexually mature adult stages. (D) Sampling locations of *I. elegans* used in the current study with inset showing the region sampled. (E) Female morph frequencies per site in 2021 and 2022. Numbers in parentheses above each bar indicate sample size. (B, C) modified from Willink et al. (2020). A = androchrome, I = *infuscans*, O = *infuscans-obsoleta*.

In addition to colour changes associated with sexual maturation, many odonates exhibit heritable (genetic) female-limited colour polymorphisms, with one female morph having male-like colouration (androchrome females) and one or two morphs expressing female-specific colouration (gynochrome females) (Blow et al., 2021; Fincke et al., 2005). Observational and experimental field studies suggest that female colour polymorphisms function to reduce the negative fitness effects of excessive male sexual harassment by disrupting males’ ability to efficiently form search images for mates (Fincke, 2004; Miller & Fincke, 1999; Takahashi et al., 2014). According to the male mimicry hypothesis, androchrome females have a negative frequency-dependent advantage in their male-like similarity, which reduces male mating harassment (Robertson, 1985). Alternatively, although not mutually exclusive, the learned mate recognition (LMR) hypothesis suggests that males form a search image for the most common female morph in the population, allowing them to better detect the most locally abundant female morph, resulting in a rare-morph advantage (Fincke, 2004). The LMR hypothesis further predicts that the ratio of correct (e.g., mature conspecific females) versus incorrect (e.g., other males, heterospecifics) male mate recognition should increase with experience (Miller & Fincke, 2004).

There is partial empirical support for both the male mimicry and LMR hypotheses and, as stated above, these two hypotheses are not mutually exclusive. In support of the LMR hypothesis, males of the damselflies *Enallagma civile* and *Ischnura elegans* show increased preference for androchrome or gynochrome females following previous exposure to these morphs (Fincke et al., 2007; Miller & Fincke, 1999; Van Gossum et al., 2001b). Further, naive *Enallagma* damselflies react sexually to both female morphs, but rarely to other males, contrary to expectations of the male mimicry hypothesis (Fincke et al., 2007). However, experiments in *Ischnura ramburi* revealed more male-male interactions in androchrome-biased settings, consistent with both mistaken mate recognition due to male mimicry and the formation of male search images for the common female phenotype (Gering, 2017). Similarly, male mating harassment in *I. elegans* increases with increasing morph density for gynochrome females, indicative of search-image formation, but not for androchrome females (Gosden & Svensson, 2009).

This growing body of ecological, evolutionary, sensory, and physiological studies, combined with the increasing availability of genomic resources (Willink et al., 2024), makes Odonata an excellent system to investigate the molecular basis of colour vision. Here, we focus on the well-studied Common Bluetail Damselfly (*Ischnura elegans*) and first characterize opsin spectral sensitivity via heterologous expression in HEK293T cells. Functional characterization of SWS and LWS rhodopsins show absorbance peaks (λ_max_) in the range of 406 – 419 nm and 531 – 548 nm, respectively. We next quantify variation in opsin mRNA expression across adult male maturation, in populations with variable female morph frequencies. Opsin expression changes are observed over adult male development, with increased expression of specific opsin types that correlate with variation in local female morph frequencies across populations. We next use visual modeling to predict the effect of shifts in visual sensitivity on female morph detection and discrimination. These results suggest that opsin expression plasticity may provide one mechanistic proximate link between male vision, morph detection/discrimination, and the resulting frequency-dependent sexual conflict, consistent with models for how these genetic polymorphisms are maintained in natural populations (Fincke, 2004; Le Rouzic et al., 2015; Svensson et al., 2005). Further experiments coupling behavioral differences in mate preferences will be interesting to link our results of ontogenetic and plastic variation in male opsin expression profiles to colour vision preferences and perception of female colour signals in this dynamic system strongly shaped by visually guided male mating behaviours.

## MATERIALS AND METHODS

### Study system

Sexually mature male *I. elegans* are monomorphic, exhibiting blue body colouration on the thorax. In contrast, female *I. elegans* are polymorphic, with sexually mature females belonging to three distinct genetically determined colour morphs (Fig. 1B) (Willink et al., 2020). Androchrome (male-like) females exhibit body colouration spectrally similar to male colouration, while gynochrome females exhibit spectrally distinct green (*infuscans*) or brown (*infuscans-obsoleta*) thorax colouration (Henze et al., 2019; Van Gossum et al., 2011). Females exhibit ontogenetic changes in thorax and abdomen colouration (Fig. 1C), with male *I. elegans* known to prefer body colouration of mature over immature females (Willink et al., 2019). Apart from colour differences, androchrome and gynochrome females also differ in resistance and tolerance to ectoparasites (Willink & Svensson, 2017) and in mating rates, resistance to mating attempts, and fecundity (Gosden & Svensson, 2009).

### Quantification of female morph frequencies between sites and field collection of males

We quantified female morph frequencies from eight sites in southern Sweden (Fig. 1D) as part of an ongoing long-term field population study of *I. elegans* (see Le Rouzic et al., 2015 and Svensson et al., 2020 for general methodology). Studied sites have both low genetic differentiation and high allelic diversity, indicating recent divergence and/or high gene flow (Abbott et al., 2008). All study populations were visited and re-visited at regular intervals (1-2 weeks) between 19 May – 1 August 2021 and 16 May – 31 July 2022. Including both immature and mature females, all populations were trimorphic except for one (Höje å 6) in 2022, which had experienced large overall population declines that year (Table S1). The frequency of each morph in the current study was calculated based on females that displayed mature adult colouration.

Male *I. elegans* were caught from sampled populations between 28 June – 13 July 2021 and 9 June – 20 July 2022 using sweep nets (Table S2). For all collected individuals, we recorded sexual maturity (“immature” vs. “mature”) by evaluating wing stiffness (Corbet, 1999). We also recorded whether males were found *in copula* with a female or singly (“couple” vs “single”). In 2022, we recorded the morph of the female that was found in copula with each collected male. In addition to immature and mature males, we also collected males that had newly emerged from the aquatic nymph stage (“tenerals”) from two field sites and from a semi-naturalistic mesocosm experiment at the Stensoffa Ecological Field Station at Lund University (Table S2). Teneral males are characterized by their extremely soft wing and body tissues and lack of body pigment. Including teneral males allowed us to assess opsin gene expression in males that have not had any significant visual experience or mated with sexually mature females. All collected males were immediately euthanized by cutting off their head using cleaned RNAse-free dissection tools, placed into individual vials filled with RNA*later* (Ambion, Inc., Austin, TX, USA) at field sites, and then stored at −20°C until RNA extraction.

### Functional expression in HEK293T cells and homology modelling

For functional expression, we selected SWS and LWS visual opsin types whose orthologs in *S. frequens* and *I. asiatica* have been shown to be expressed in the adult ventral compound eye (SWb1, SWb2, LWA2, LWF1 – F4) and LWE1 to test for function of an opsin whose ortholog in *I. asiatica* is primarily expressed in ocelli (Futahashi et al., 2015). Homology modelling was performed for the SWb1 and LWF1 opsin amino acid sequences, aligned against the invertebrate jumping spider rhodopsin crystal structure (PDB 6i9k) (Varma et al., 2019). Full methods are presented in supplemental methods.

### Quantitative PCR analysis of *I. elegans* opsin expression levels

We quantified whole head opsin mRNA expression via quantitative PCR (see supplemental methods) in N = 87 male *I. elegans* (8 teneral, 36 immatures, 43 matures; Table S2) for UV, SWb1, SWb2, LWA2, LWF1-F4 and LWE1 opsins. Hereafter, we use abbreviated names for the opsins SWb1, SWb2, LWA2, and LWE1, which have been shortened to SW1, SW2, LWA, and LWE, respectively.

Expression of opsin genes was calculated relative to two housekeeping genes and calibrated against the UV opsin gene using the (1 + E)^−ΔΔCT^ method where E equals the PCR efficiency for each primer (Livak & Schmittgen, 2001; Pfaffl, 2001). We also calculated the proportion opsin expression relative to total opsin expression for opsins likely expressed in the main compound eye in other odonatans by dividing relative expression for each individual opsin by the sum of total opsin expression. Results of relative opsin expression have been shown to indicate differential regulation of opsin genes, while proportional measures of opsin expression allow to make inferences about colour vision (Fuller & Claricoates, 2011).

### Visual modelling of female morph detection and discrimination

To assess how observed differences in LWS opsin expression might impact detection or discrimination of female morphs, we focused on the two most abundant female morphs (androchrome and *infuscans*) and calculated just noticeable differences (JNDs) for three parsimonious variations of Odonate visual systems. The first modelled visual system follows ERG spectral sensitivity values from *Ischnura heterosticta* with SWS λ_max_ = 450 nm and LWS λ_max_ = 525 nm (Huang et al., 2014). For the second and third visual models, based on *I. elegans* opsin spectral sensitivity, we calculated JNDs with SWS λ_max_ = 413 nm, together with LWS λ_max_ = 531 nm (model 2) or 543 nm (model 3). The LWS spectral sensitivities correspond to maximal opsin absorbance spectra for *I. elegans* LWF1 and LWF2, for which both relative and proportional gene expression decreased and increased, respectively, in mature versus immature males (see results). All models included a fixed UVS spectral sensitivity (λ_max_ =360 nm), consistent with published electroretinogram (ERG) data in other Odonate species (Futahashi et al., 2015; Huang et al., 2014). Equations used in visual model calculations are presented in supplemental methods.

### Statistical analysis

We performed a Multivariate Analysis of Variance (MANOVA) to assess the effect of maturity on log relative opsin gene expression of all eight opsin genes for male *I. elegans.* Log relative opsin expression for each individual male was averaged between replicates to maintain the assumption of independence.

We used linear mixed modeling to test the effects of population morph frequency, maturity stage, and opsin type on log relative and proportion opsin expression in male *I. elegans*, with site, male ID, year, and qPCR replicate as possible random effects. Model selection followed methods in Zuur et al. (2009) to identify the random and fixed effects included in each model. In total, we ran three main models. **Model 1** assessed changes in relative opsin expression for the main effects of maturity stage and opsin type, including opsins likely expressed in both the compound eyes and ocelli. **Model 2** assessed changes in proportion opsin expression for opsins with likely expression in only the compound eye, with the main effects of maturity stage and opsin type. **Model 3** tested the effects of maturity stage, opsin type, population morph frequency and its squared component on relative opsin expression. Detailed modelling methods are presented in supplemental methods.

We then assessed whether there were differences in opsin expression for males captured *in copula* with androchrome or *infuscans* females for males captured in 2022. However, model selection suggested no significant effect of the female morph that a male was found mating with on relative opsin expression, possibly due to relatively small sample sizes. We therefore do not discuss these results in the main text (see supplemental methods).

## RESULTS

### Female morph frequencies

The frequencies of the three mature female morphs differed between sites and years (Fig. 1E-F). Androchrome females (A) were the majority female morph at all sites, and their frequency ranged from 57.1 – 85.7% (mean ± sd: 72.6 ± 9.7%) in 2021 and 65.9 – 83.9% (72.3 ± 5.8%) in 2022. The *infuscans* female morph (I) was the next most common morph, ranging from 14.3 – 39.1% (24.1 ± 8.8%) in 2021 and 16.1 – 30.7 (24.9 ± 5.3%) in 2022. Finally, the third morph (*infuscans-obsoleta*, O) were rare, making up only between 0.00 and 5.5% (3.3 ± 1.8%) of mature females in 2021 and between 0.00 and 7.8% (2.8 ± 2.8%) in 2022. Morph fluctuations across years is common in this species, although in many populations morph frequencies are relatively stable over time and across many generations, as a result of negative frequency-dependent selection (Le Rouzic et al., 2015; Svensson & Abbott, 2005). Accordingly, only one of these eight populations differed significantly in morph frequencies between these two years (Vombs Vattenverk). This population had a 1.7-fold decrease in the frequency of I-females and a 10-fold decrease in the frequency of O-females relative to A-females between the two years (see Table S3 for statistical comparisons for all sites).

### Opsin spectral sensitivity determined using HEK293 cell heterologous expression

We found that the SWS opsins SWb1 and SWb2 absorb maximally at λ_max_ of 406 and 419 nm (Fig. 2A-B). The LWS opsin types display maximal sensitivity between 530 and 550 nm with the following λ_max_: LWA2 = 548 nm, LWF1 = 531 nm, LWF2 = 543 nm, LWF3 = 545 nm, LWF4 = 541 nm, LWE1 = 533 nm (Fig. 2C-H). Spectral sensitivity functions of SWS and LWS opsins overlap with spectral reflectance of female thorax coloration of androchrome and *infuscans* females (Fig. 3).

**Figure 2.**
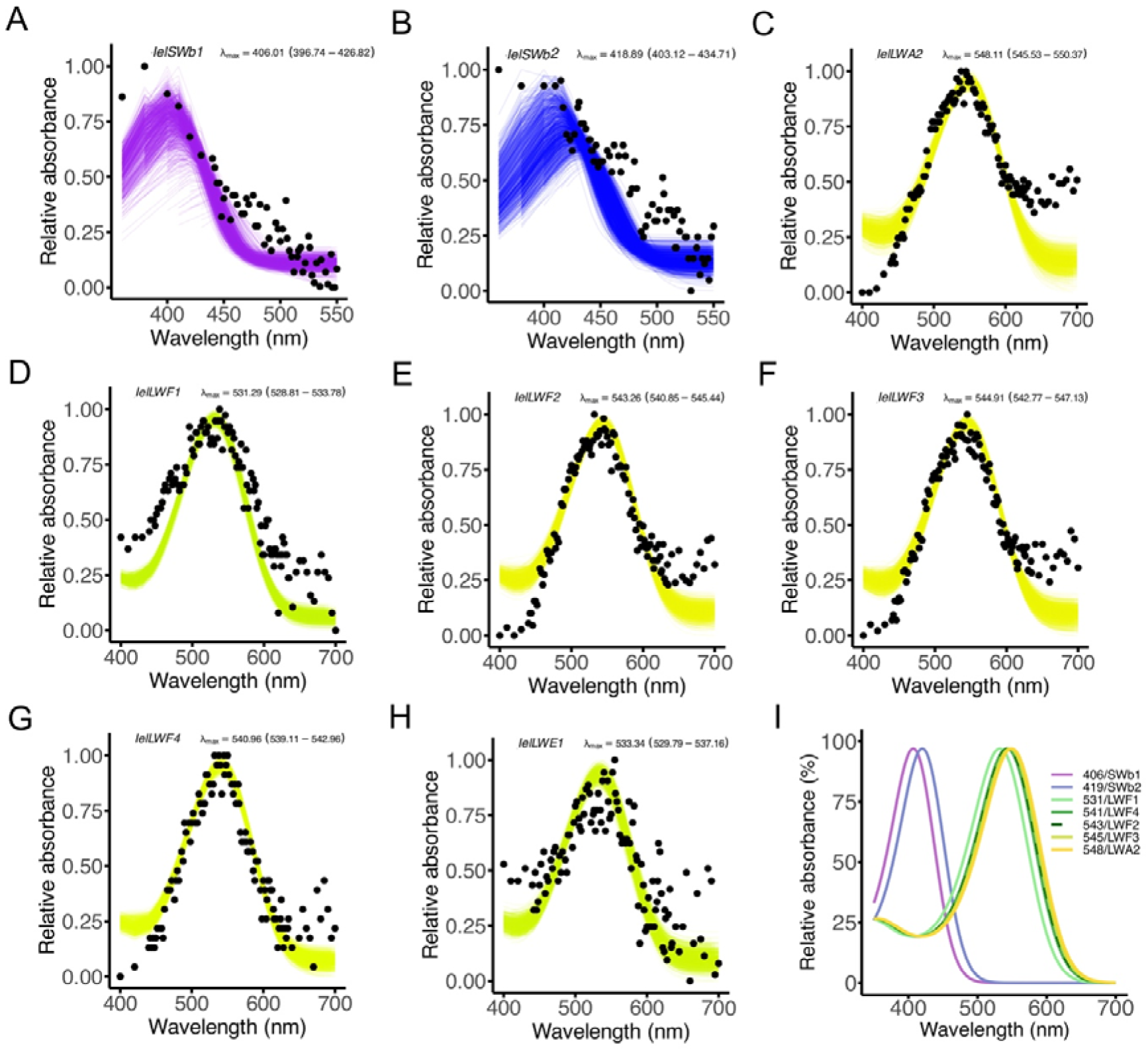
Functional expression of *Ischnura elegans* short-wavelength (SWS) and long-wavelength (LWS) opsins. Ultraviolet-visible (UV-VIS) absorbance analyses of dark-adapted rhodopsin visual pigments reconstituted and purified from HEK293T cell cultures in the dark in the presence of 11-*cis*-retinal. The black dots represent mean absorbances at a given wavelength. (A) SWb1 (n = 7), (B) SWb2 (n = 3), (C) LWA2 (n = 2), (D) LWF1 (n = 2), I LWF2 (n = 5), (F) LWF3 (n = 6), (G) LWF4 (n = 1), and opsins expressed in the ocelli: (H) LWE1 (n = 6), where n is the number of measurements of protein aliquots with active rhodopsin complexes. (I) Combined absorbances for SWS and LWS opsins, excluding LWE1, which is expressed in the ocelli in other odonates. Absorbance at 380 nm in A and B is due to residual unbound *cis*-retinal. Relative absorbance data are fitted to a visual template with polynomial function analyses computed in R to obtain the best estimates following 1000 bootstrap analysis of lambda max for each rhodopsin (Liénard et al., 2022). Confidence intervals are indicated in parentheses.

**Figure 3.**
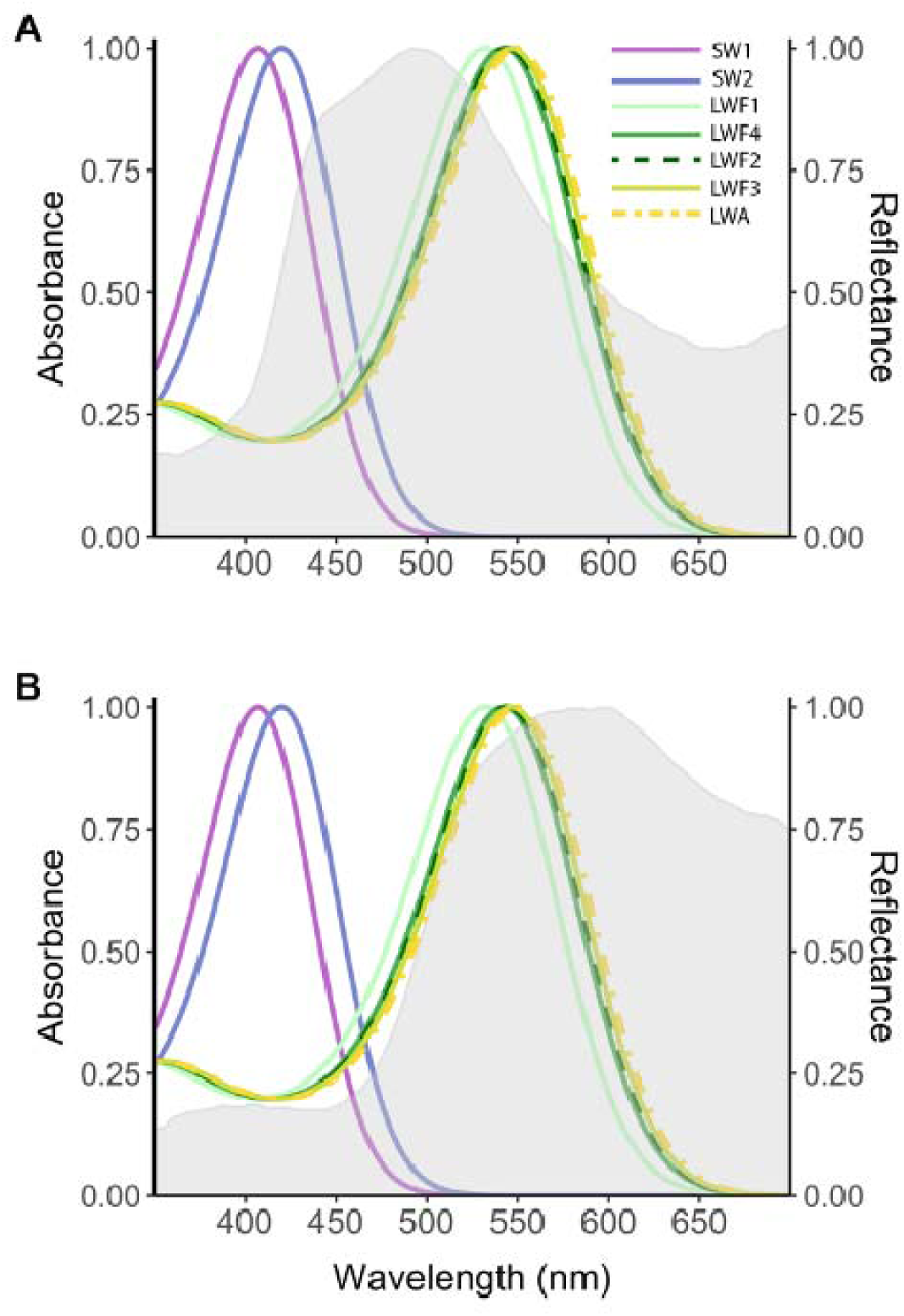
Visual opsin sensitivity and female body reflectance. Absorbance spectra of *I. elegans* SWS and LWS opsins characterized in this study are represented by coloured visual fit curves and overlapped with (A) androchrome and (B) *infuscans* female body reflectance (shaded gray areas). Spectral sensitivity functions correspond to differences in achromatic (brightness) and chromatic cues (body colour reflectance), assuming opponency between SWS and LWS opsins, of androchrome and *infuscans* females. SW1, SW2 and LWA labels correspond to SWb1, SWb2 and LWA2 opsins, respectively. LWE1, which is primarily expressed in the ocelli of other odonates, is not included. Body reflectance data from Henze et al. (2019).

We ran homology modeling of SWb1 and LWF1 opsins against the spider rhodopsin (PDB: 6i9k) and mapped sites interacting with the chromophore in PyMOL. Of the 71 amino acid substitutions between SWb1 and SWb2 (81.8% aa identity), we identified 23 sites predicted to be within 5Å of the binding pocket (Table S4). Among these, Y136 (TM3) and Y294 (TM6) residues located at the top and bottom of the binding pocket, respectively, are substituted by phenylalanine (F136 and F294) in SWb2 (Fig. S1A). For LWS opsins, which share between 88.9 - 91.2% aa identity (i.e., 33 - 42 variant residues), we obtained the predicted LWF1 opsin structure based on 6i9k and mapped 24 sites predicted to interact with the cis-retinal, all of which are conserved for all four LWF opsins (Table S4; Fig. S1B). The LWF2 protein shares 89.9-90.5% identity with LWF1-F4, translating to 36 to 38 residue substitutions, and 67.4% identity (123 amino acid residue differences) with LWA2 (Fig. S3B). LWA2 is more divergent in sequence, 67.5 - 68.7% aa identity with 119 - 124 aa differences with LWF opsins. Of the 24 chromophore-interacting residues, its binding pocket exhibits two potential spectral variant residues with LWF1 (A131G in TM1 and S319A in TM7).

### Opsin expression in teneral, immature, and mature males

We compared the expression levels of 2 SWS and 6 LWS opsins, relative to UV opsin expression, across adult male developmental stages in *I. elegans* from eight sites. Our results from the MANOVA showed a significant effect of male maturity stage on overall opsin gene expression (F_16,146_ = 3.2, *p* < 0.0001; Fig. 4). To further assess how relative and proportion opsin expression differed between teneral, immature, and mature male *I. elegans*, we performed mixed modeling with maturity stage, opsin type, and their interaction as main effects.

**Figure 4.**
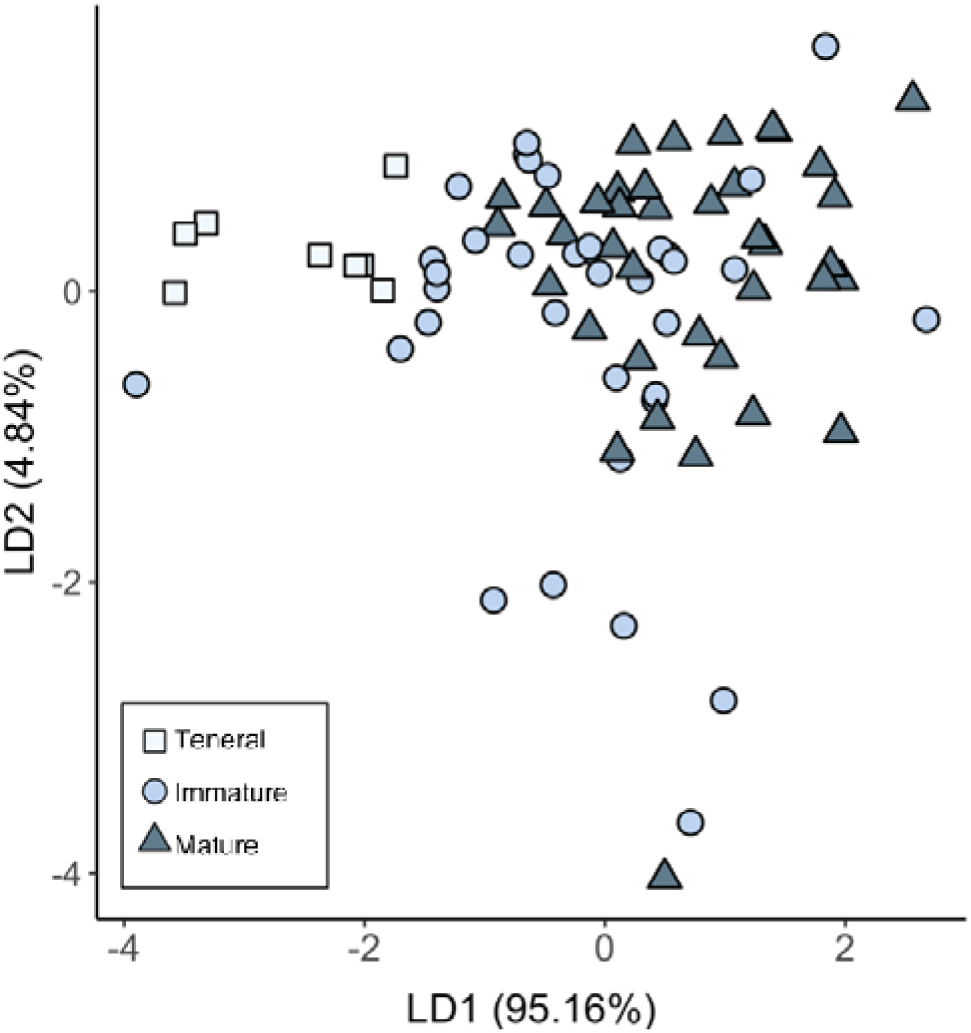
Linear discriminate analysis showing the relationship between relative expression of eight opsins expressed in teneral, immature, and mature male *I. elegans.* The percent of variation described by each linear discriminate factors (LD1 and LD2) are shown in parenthesis. Distance between points across axes indicates differences in overall relative opsin expression between maturity stages, with results showing the largest differences between teneral and mature males, with immature males intermediate to the two groups.

For relative opsin expression (model 1), we found a significant interaction between maturity stage and opsin (Table 1, Tables S5 – S6) with increasing differences in relative expression from teneral males (Fig. 5A) to immature (Fig. 5B) and mature (Fig. 5C) males. For teneral males, highest relative opsin expression was for LWF4 (Fig. 5A), while LWF2 has the highest relative expression for immature and mature males (Fig. 5B, C). Results also show that, for immature and mature males, the opsin LWE has significantly lower relative expression relative to other opsin types, consistent with expression levels observed in S*. frequens* and *I. asiatica* (Futahashi et al., 2015).

**Figure 5.**
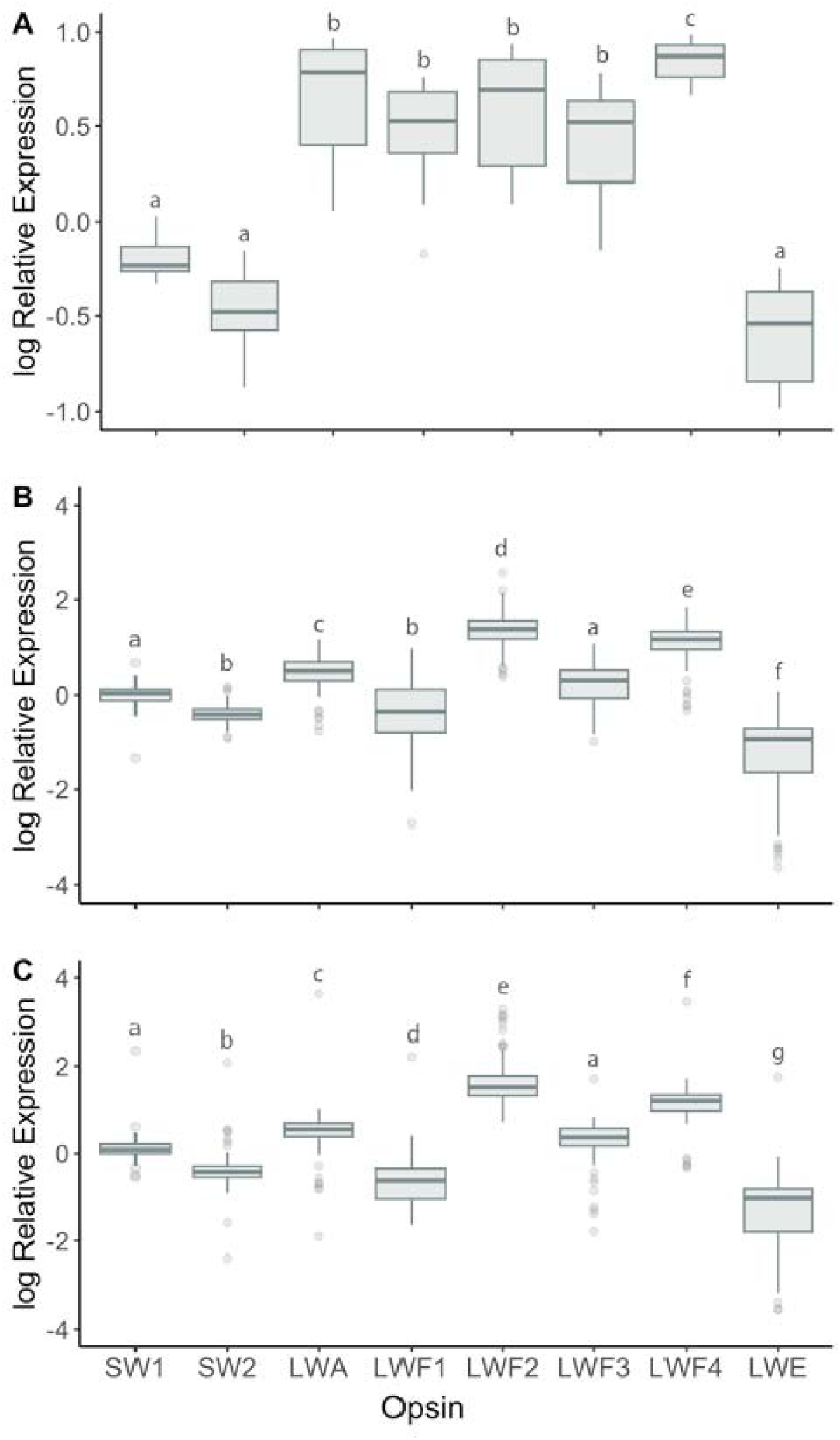
Relative opsin expression for (A) teneral, (B) immature, and (C) mature male *I. elegans* across 8 sites. Bars in box and whisker plots show medians, boxes indicate upper and lower quartiles, whiskers show sample minima and maxima, and open circles show outliers (representing 5.2 and 6.4 % for immature and mature treatments respectively). Expression data are normalized against housekeeping genes and calibrated against UVS opsin expression levels. Letters above each bar show significant differences between opsin types within each maturity level following Tukey corrections for multiple comparisons. Test statistics and p-values for all comparisons are presented in Table S5. Single population analyses are presented in Fig. S2.

**Table 1.** Final model and model output for model 1, showing the effect of maturity stage and opsin type on log relative opsin expression in teneral, immature, and mature male *I. elegans*. Significant differences are shown in italics.

| log Relative Opsin Expression ~ Maturity × Opsin + (1\|ID) + (1\|Replicate) + (1\|Year) |  |  |  |
| --- | --- | --- | --- |
|  | df | F | P |
| Maturity | 2 | 0.2 | 0.82 |
| Opsin | 7 | 292.3 | < 0.0001 |
| Maturity × Opsin | 14 | 12.3 | < 0.0001 |

Comparing teneral, immature, and mature males, results showed that expression changes over male maturation are not parallel across opsins, but that there were opsin-specific changes (Fig. 6). Increases in relative expression occurred for the SW1 (teneral – mature: t = −2.7, df = 396, *p* = 0.02) and LWF2 opsins (teneral – immature: t = −5.1 df = 401, *p* < 0.0001; teneral – mature: t = −6.9, df = 396, *p* < 0.0001; immature – mature: t = −3.0, df = 419, *p* = 0.009). Conversely, decreases in relative expression occurred for the LWF1 (teneral – immature: t = 4.7, df = 402, *p* < 0.0001; teneral – mature: t = 6.1, df = 399, *p* < 0.0001; immature – mature: t = 2.4, df = 433, *p* = 0.04) and LWE opsins (teneral – immature: t = 3.4, df = 401, *p* = 0.002; teneral – mature: t = 3.9, df = 400, *p* = 0.0004). For results for each independent population, see supplemental materials (Table S7, Fig. S2).

**Figure 6.**
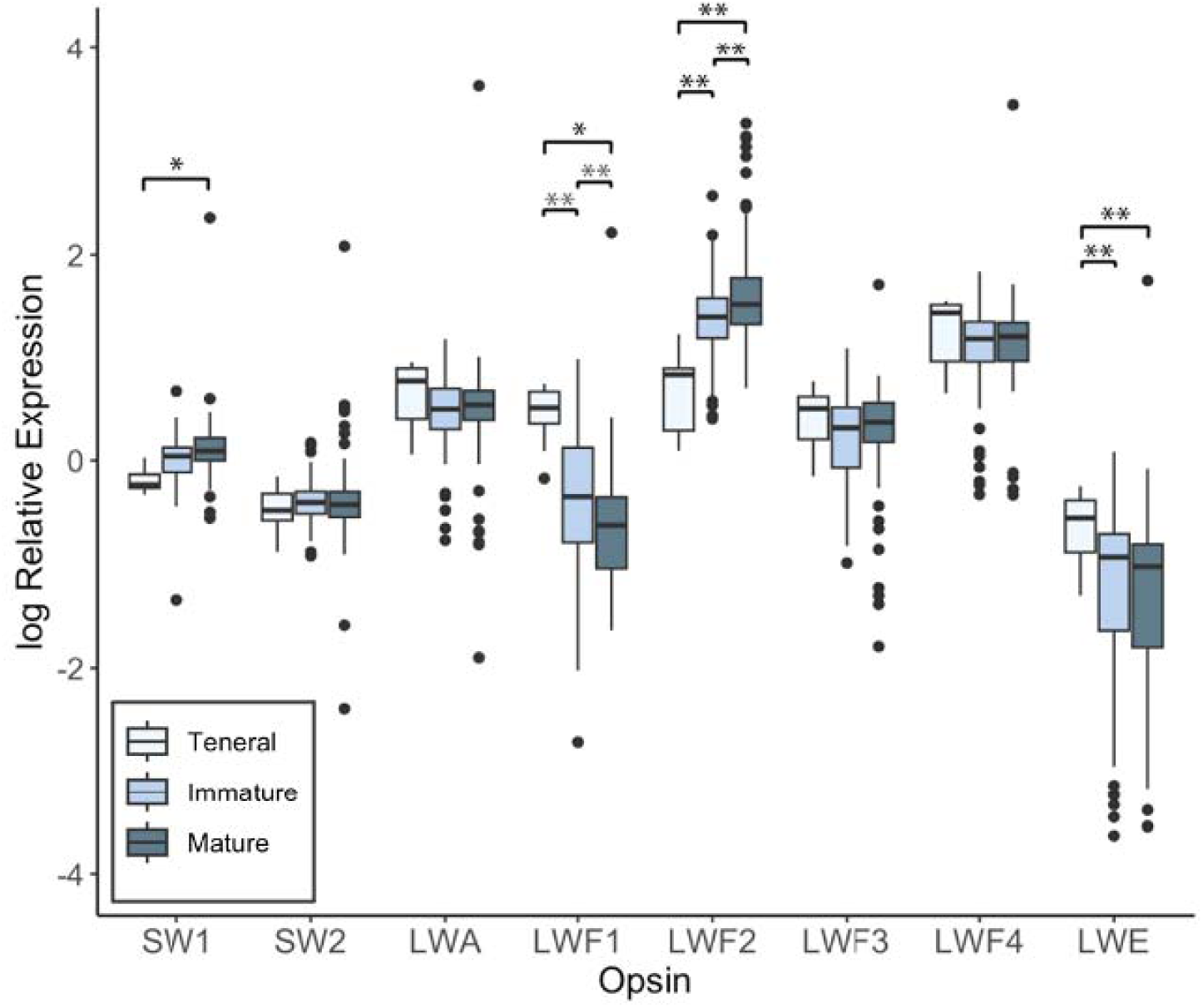
Relative male opsin expression across eight *I. elegans* populations for seven opsins expressed in the compound eye (SW1 – LWF4) and one low-expressed opsin likely expressed in the ocelli (LWE). Teneral males are shown in light blue, immature adult males in medium blue, and mature males in dark blue. Bars in box and whisker plots show medians, boxes indicate upper and lower quartiles, whiskers show sample minima and maxima, and open circles show outliers (representing 5.2 and 6.4 % for immature and mature treatments respectively). Expression data are normalized against housekeeping genes and calibrated against UVS opsin expression levels. Single asterisks indicate comparisons that are significant at alpha < 0.05 and double asterisks represent comparisons that are significant at alpha < 0.01 following Tukey corrections for multiple comparisons within opsin types. Test statistics and p-values for all comparisons are presented in Table S6; single population analyses are presented in Fig. S2.

When analyzing proportional opsin expression (model 2), we again found a significant interaction between maturity stages and opsin type (Table 2). For all LWS opsin types, with the exception of LWF1, we found a significant difference between teneral males and both immature and mature males, but not between immature and mature males (Table S8). For LWF1, there was a significant difference between all maturity stages tested (teneral – immature: t = 12.4, df = 84, *p* < 0.001; teneral – mature: t = 15.2, df = 84, *p* < 0.001; immature – mature: t = 4.4, df = 84, *p* < 0.001). Relative and proportional opsin expression were thus largely consistent, with a decrease in LWF1 expression and increase in LWF2 expression over maturity, although differences in proportional LWF2 expression were not significant between immature and mature males (t = 2.1, df = 84, *p* = 0.10), as with relative expression (Fig. 7). LWF4 expression, which did not significantly differ in relative expression over development (Fig. 6), made up the largest proportion expressed for teneral males (0.523 ± 0.01). In immature and mature males, LWF2 made up the largest proportion opsin expressed, accounting for 0.524 ± 0.03 and 0.599 ± 0.02 of expression, respectively (Fig. 7).

**Figure 7.**
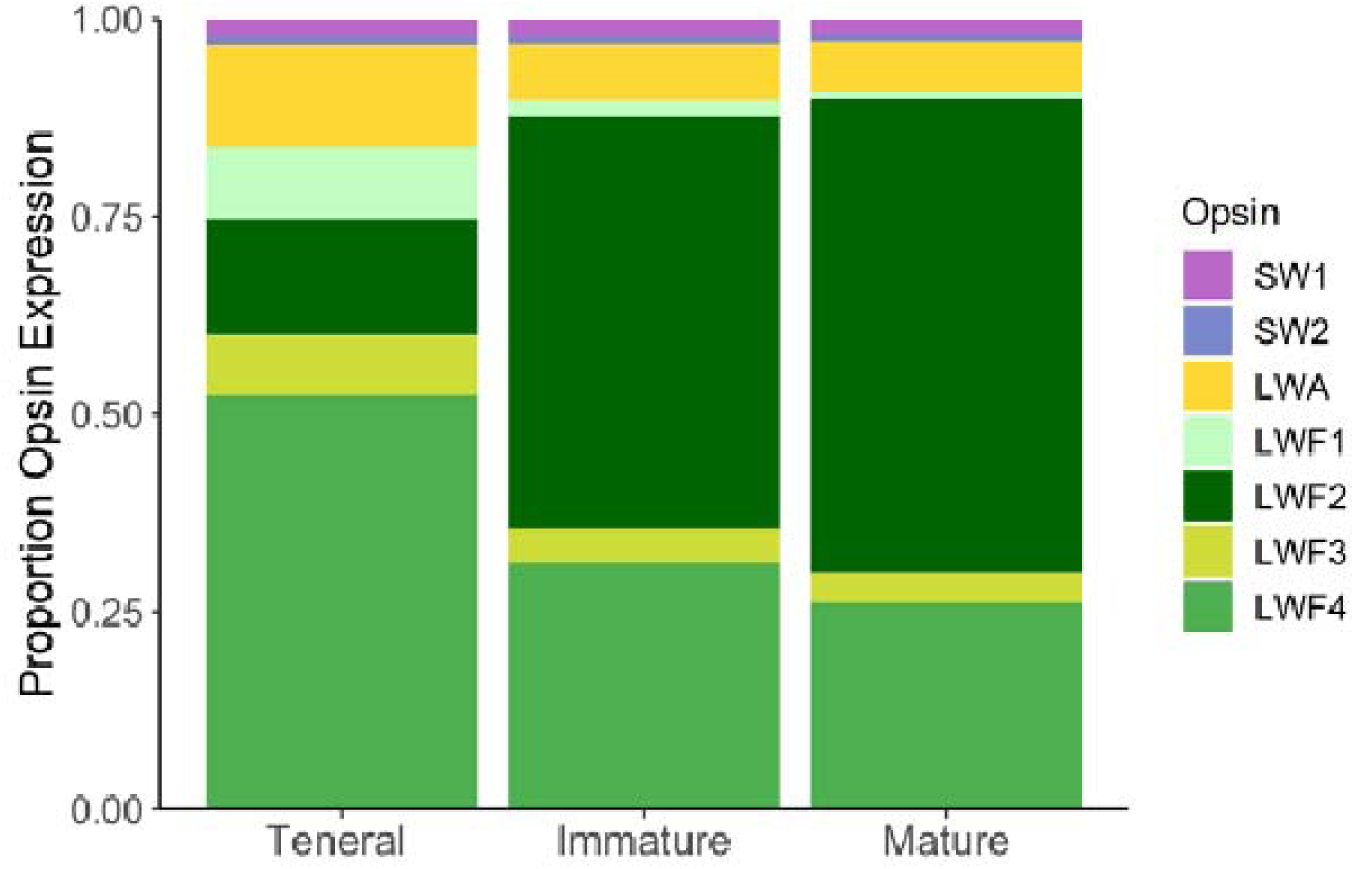
Average proportion expression of visual opsins in teneral, immature, and mature male *I. elegans*. Proportion expression for 2 short-wavelength sensitive (SWS) and five long-wavelength sensitive (LWS) opsins.

**Table 2.** Final model and model output for model 2, showing the effect of maturity stage and opsin type on proportion opsin expression in teneral, immature, and mature male *I. elegans*. Significant differences are shown in italics.

| Proportion Opsin Expression ~ Maturity × Opsin + (1\|ID) |  |  |  |
| --- | --- | --- | --- |
| | df | $\chi^2$ | P |
| Maturity | 2 | 20.7 | < 0.0001 |
| Opsin | 6 | 8877.3 | < 0.0001 |
| Maturity × Opsin | 12 | 437.5 | < 0.0001 |

### Opsin expression across populations with different female morph frequencies

Comparing variation in opsin expression in immature and mature males including the effect of local female morph frequency (model 3), we find a significant interaction between opsin type, maturity stage, and the squared component of androchrome frequency (Table 3). Plotting log relative expression for each opsin type revealed a quadratic relationship between LWF2 expression and andromorph frequency in mature, but not in immature, males. We observed the highest LWF2 expression in populations with the lowest and highest proportion of androchrome females. In contrast, androchrome frequency had little effect on relative expression for immature or mature males for the other opsins tested (Fig. 8).

**Table 3.** Final model and model output for model 3, showing the effect of the proportion of androchrome females in a population, opsin type, and maturity stage on log relative opsin expression for immature and mature male *I. elegans*. Significant differences are shown in italics.

| log Relative Opsin Expression ~ Opsin × Maturity × (Prop. Androchrome + Prop. Androchrome^2) + (1\|ID) + (1\|Replicate) + (1\|Year) |  |  |  |
| --- | --- | --- | --- |
|  | df | F | P |
| Opsin | 7 | 18.7 | < 0.0001 |
| Maturity | 1 | 1.3 | 0.27 |
| Prop. Androchrome | 1 | 0.2 | 0.66 |
| Prop. Androchrome^2 | 1 | 0.2 | 0.67 |
| Opsin × Maturity | 7 | 3.1 | < 0.01 |
| Opsin × Prop. Androchrome | 7 | 17.1 | < 0.0001 |
| Opsin × Prop. Androchrome^2 | 7 | 17.3 | < 0.0001 |
| Maturity × Prop. Androchrome | 1 | 1.3 | 0.26 |
| Maturity × Prop. Androchrome^2 | 1 | 1.3 | 0.25 |
| Opsin × Maturity × Prop. Androchrome | 7 | 3.3 | < 0.01 |
| Opsin × Maturity × Prop. Androchrome^2 | 7 | 3.6 | < 0.001 |

**Figure 8.**
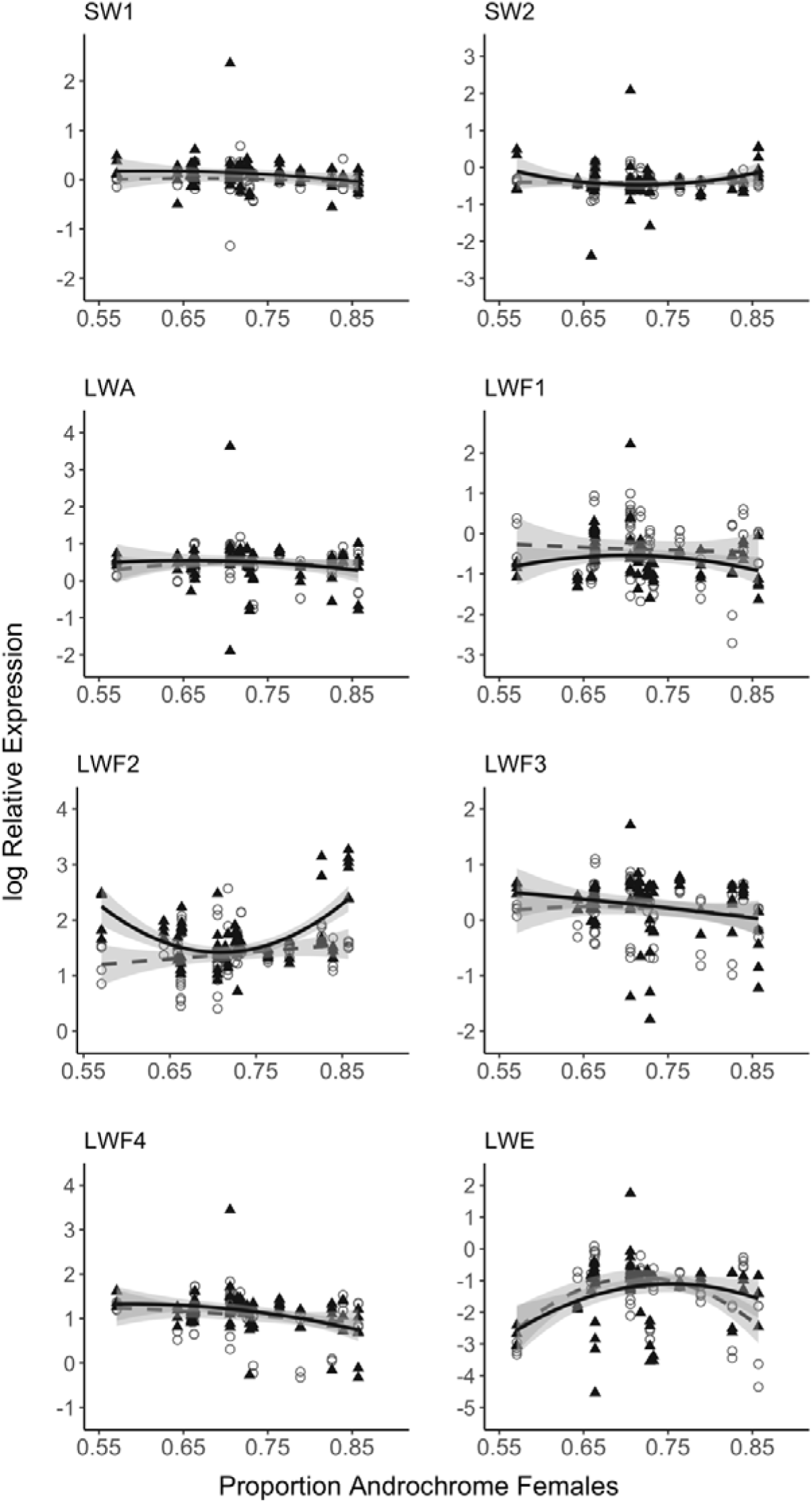
Opsin expression changes relative to proportions of androchrome females. Log relative opsin expression in immature (open circles, dashed line) and mature (black triangle, solid line) male *I. elegans* from populations with different proportions of androchrome females. Grey shading around trend lines indicates 95% confidence intervals.

### Visual modelling of female morph detection and discrimination

Comparing the results of visual models using ERG data from *I. heterosticta*, *I. elegans* LWF1 opsin sensitivity, and *I. elegans* LWF2 opsin sensitivity, we found that the LWF2 visual system had larger just-noticeable differences (JNDs) for discriminating androchrome and *infuscans* color morphs (ΔJND_LWF2-ERG_ = 0.14, ΔJND_LWF2-LWF1_ = 0.41) and for detecting androchrome females against brown and green vegetation (brown: ΔJND_LWF2-ERG_ = 0.53, ΔJND_LWF2-LWF1_ = 0.56; green: ΔJND_LWF2-ERG_ = 0.14, ΔJND_LWF2-LWF1_ = 0.36). Modelling with parameters from the *I. heterosticta* ERG visual system led to larger JNDs for detection of *infuscans* females against brown vegetation (ΔJND_LWF1-ERG_ = −0.16, ΔJND_LWF2-ERG_ = −0.17) (Table S9). The three tested visual systems performed similarly well for discrimination of *infuscans* females against green vegetation (ΔJND_LWF2-ERG_ = 0.02, ΔJND_LWF2-LWF1_ = 0.07, ΔJND_LWF1-ERG_ = −0.05).

## DISCUSSION

Vision is an important sensory modality in Odonata and the prevalence of female-limited colour polymorphisms makes the damselfly genus *Ischnura* a powerful model system to study how and why polymorphisms emerge and persist over both micro- and macroevolutionary time scales (Blow et al., 2021; Le Rouzic et al., 2015; Willink et al., 2024). Here, we combined ecological, molecular, and functional approaches to investigate opsin gene expression in male *I. elegans* of different maturation stages and across populations with varying female morph frequencies. Our measures of *in vitro* opsin absorbance evidence that the photosensitive opsin receptors forming the peripheral visual system of *I. elegans* perceive wavelengths that overlap with reflectance intensity (brightness) and chromatic body colouration cues of the two most abundant female morphs, namely androchrome and *infuscans*, from field sites (Fig. 3). Further, recurrent directional changes in opsin expression over male ontogeny, particularly during sexual maturation (Figs. 4 - 7), as well as changing male gene expression profiles in response to local female morph frequencies (Fig. 8) suggest a role for opsin expression plasticity in the *Ischnura* system.

Vision is used for many tasks in Odonates, and ecological factors might select for additional differences that may not necessarily be attributable to conspecific search and morph discrimination. The spatial organization of yet unknown ommatidial types possibly expressing a subset of the multiple SWS and LWS opsins across the retina may also influence how male *I. elegans* perceive conspecific body cues. However, *I. elegans* is equipped with numerous functional SWS and LWS visual opsins activated maximally by wavelengths of light highly reflected by female morphs (Fig. 3). In light of known behavioural male preference for female colour morphs and dynamically changing morph frequencies (Van Gossum et al., 1999, 2001b), opsin-based spectral sensitivity across short and long wavelengths likely contributes to visual detection and discrimination of local female morphs.

### Overlapping opsin spectral sensitivities likely confer broad wavelength discrimination

A total of 12 expressed visual opsin genes have been identified in the damselfly *Ischnura asiatica* (Futahashi et al., 2015), a gene repertoire that we find conserved in the genome of *I. elegans* (Fig. S3A). From the *I. elegans* opsin gene set, we quantitatively assessed whole head adult expression levels of 2 SWS (SWb1, SWb2) and 5 LWS (LWA, LWF1-F4) opsins for which orthologs in *I. asiatica* and *Sympetrum frequens* are primarily expressed in the ventral eye (Futahashi et al 2015), a region implicated in conspecific search (Lancer et al., 2020). For comparison, we also quantified expression levels of *I. elegans* LWE1 opsin, which is restricted to ocelli in other odonates (Futahashi et al 2015). The *I. elegans* LWE1 opsin, which is sensitive to green wavelength light (Fig. 2), consistently exhibited the lowest monitored expression levels across opsin types (Fig. 3), suggesting a restricted ocelli-specific expression pattern and function. In contrast, SWS and the other LWS opsins exhibited higher baseline expression levels compared to LWE1, as observed in *I. asiatica,* therefore supporting a role as visual opsins in the main compound eye.

The SWS opsins confer sensitivity to light in the violet-blue spectrum (λ_max_ = 406 nm and 419 nm) and the five visual LWS opsins show broad sensitivity to green wavelengths of light, with λ_max_ between 530 nm and 540 – 550 nm (Fig. 2). This forms a key component defining the genetic basis of light perception and functionality of multiple opsin duplicates, as part of several complementary mechanisms that certainly contribute to modulate visual sensitivity and capture a remarkably broad light spectrum from ultraviolet to long-wavelengths above 600 nm. While achromatic vision of intensity related cues can be directly modulated by light capture of single opsins (Kelber, 2005), chromatic discrimination involves at least two distinct but overlapping sensitivities in a given ommatidial unit (Kelber & Osorio, 2010; Lind et al., 2017). ERG recordings from other odonates show broad sensitivity of the green photoreceptor type which may indicate the presence of multiple opsin genes expressed within a single photoreceptor (Futahashi, 2016; Huang et al., 2014). Irrespective of how many LWS opsins may be expressed in a single photoreceptor or ommatidium, overlap between short- (UV and violet-blue) and at least one long-wavelength opsin sensitivities could provide fine-tuned trichromatic discrimination capacity across a broad spectral range in *I. elegans* (Figs. 2, 3), with additional filtering mechanisms common in arthropods potentially further increasing spectral channels (Cronin et al., 2001; Stavenga & Arikawa, 2011).

Our results show a high degree of overlap in the recorded sensitivities of the LWS opsins, which suggest that changes in expression of any single opsin might not have a large impact on spectral sensitivity if several LWS opsins are co-expressed. This also suggests that there may be no opponency unless combining LWS opsins with 10-20 nm differences between peak spectral sensitivity (Kelber et al., 2003). However, substantial increases and decreases in opsin expression levels for opsin types with λ_max_ differences greater than 10 nm (e.g., LWF1 λ_max_ = 531 nm, and LWF2 λ_max_ = 543 nm), may affect the underlying photoreceptor spectral visual sensitivity if these specific opsins are also spatially segregated across the retina.

Spatial differences in opsin expression patterns have been identified in other insects (Rister & Desplan, 2011) including other species of Odonata (Futahashi et al., 2015; Labhart & Nilsson, 1995), which suggests that *I. elegans* LWS opsin duplicates could have regionalized expression patterns throughout the retina. For example, the eyes of the dragonfly *Hemicordulia tau* seem to have three short-wavelength sensitive zones (i.e., the dorsal acute zone, the dorsal rim area, and the frontal acute zone) and display ventral spectral band patterning maximally sensitive to either short-, middle-, or long-wavelengths (Lancer et al., 2020). Likewise, ERG recordings from the dragonfly *S. frequens* show regional differences in spectral sensitivity coinciding with dorso-ventral opsin expression levels and broad green spectral sensitivity (Futahashi et al., 2015). Future analysis of LWS opsin spatial distribution via gene-specific antibody staining or *in situ* hybridization, along with *in vivo* ERG quantification of visual sensitivity, will be valuable next steps to determine how opsin spatial distribution and co-expression patterns may contribute to finely tune visual spectral sensitivity in *I. elegans*.

### Candidate spectral tuning sites between SWS and LWS opsin duplicates

Homology modeling indicated that the residues predicted to interact with the retinal chromophore are conserved between all tested LWF opsins. It is therefore possible that variant residues at positions outside the canonical binding pocket play a role in fine spectral tuning, causing the observed narrow spectral shifts.

For SWS opsins, we mapped two strong candidate spectral tuning sites at positions 136 and 294 between *I. elegans* duplicate SWb1 (λ_max_ 406 nm) and SWb2 (λ_max_ 419 nm) opsins (Fig. S1). A comparison of duplicate SWb opsins across five *Ischnura* species (see SI dataset 2), *I. cervula*, *I. verticalis*, *I. hastata* and *I. asiatica,* shows that Y294F (H6) is present only in *I. cervula* whereas Y136F (H3) is conserved across all species. Although we have not tested the effects on these mutations in the present study, repeated homologous Y/F changes have been functionally shown to cause convergent spectral shifts in SWS insect opsins (Frentiu et al., 2015; Liénard et al., 2021; Wakakuwa et al., 2010). Accumulated evidence for the role of Y to F substitutions in providing longer wavelength-sensitive SWS invertebrate opsins is consistent with the gain of F136 in *I. elegans* SWSb2, possibly underlying the change in paralog SWS sensitivity (Fig. 2). The conserved Y136F substitution presumably also influences spectral tuning across other *Ischnura* SWb1 and SWb2 opsin paralogs, although possibly to varying degrees, as species-specific variation in spectral sensitivity can be expected between orthologous native opsins, as observed with the orthologous opsins of *S. frequens*: LWA2 (λ_max_ 557 nm) and LWD1 (λ_max_ 542 nm) (Liénard et al., 2022) compared to the *Ischnura* orthologs (LWA2 λ_max_ 548 nm; LWD1 λ_max_ 541 nm) characterized here.

### Opsin expression plasticity as a potential mechanism underlying morph detection and discrimination

In this system with rapidly changing female morph frequencies, males are expected to rapidly develop plastic search images, which may be shaped by local frequencies of female morphs and their intrinsic fecundities (Fincke, 2004; Gosden & Svensson, 2009; Verzijden et al., 2012). The visual system and peripheral opsin receptors are known major evolutionary targets (Dungan & Chang, 2022; Hauzman et al., 2021; Macias-Muñoz et al., 2016; Schott et al., 2024) with the potential to fine tune wavelength light capture and colour vision, in particular in altering opsin gene evolution, function and expression. There is also increasing empirical evidence for a key role of phenotypic plasticity in evolution, including in shaping the development of both male and female mate preferences (Verzijden et al., 2012; West-Eberhard, 2003). Such learned mate preferences may impact sexual selection, sexual conflict, the maintenance of sexually selected polymorphisms, population divergence, and ultimately speciation (Svensson et al., 2010; Verzijden et al., 2012; Westerman et al., 2012).

The damselfly *I. elegans* is characterized by visually guided mate-searching males who are exposed to female colour polymorphisms and a continuum of androchrome female frequencies across local populations (Fig. 1). This system allowed us to explore male visual system response dynamics quantitatively in natural field settings. Quantification of opsin mRNA levels throughout adult *I. elegans* maturation at eight sampling sites over two flying seasons revealed consistent plasticity for specific opsin genes over adult male development. In particular, we identified opsin-specific directional changes in relative expression for LWF1 and LWF2 and proportional changes in the same opsin genes, in addition to the other LWS opsin types tested, over male ontogeny across several populations (Figs. 4 - 7). This suggests that males might experience age-specific selection pressures on these vision genes. Successful visual mate identification has also been predicted to vary in response to local female morph abundances and correlate with increased sexual experience (Fincke, 2004; Miller & Fincke, 2004). Since opsin expression profiles of adult males were consistent with a gradual increase (i.e., LWF2) and decrease (i.e., LWF1) in opsins with distinct spectral sensitivities, these changes may contribute, at least partly, in fine-tuning the male visual system for mate detection and mate discrimination during sexual maturation.

Moreover, sexually immature teneral males are inexperienced with visually guided behaviors compared to immature and mature adult males. In line with this, we found that opsin gene expression levels, for SWS and LWS visual opsin types, are more homogenous in teneral whole head tissues, whereas variation in relative expression increased in immature and mature male *I. elegans* (Fig. 5). Major directional expression changes were found at the metapopulation level, including changes that reliably depend on maturity stage, and target opsin type, with notable decreases in LWF1 and increases in LWF2 expression (Fig. 6). The relationship between LWF1 and LWF2 expression appeared to be conserved across all eight sampled populations, and statistical signal is recovered at the population level for LWF1 and LWF2 in several of the individual sites (Fig. S2), despite lower sampling power. Taken together, these differences in relative expression are consistent with opsin-specific regulation during ontogenic development and male sexual maturation (Fig. 6), suggesting that vision changes in immature and mature adult males compared to teneral males.

Proportional opsin expression also differed between developmental stages, with significantly more abundant LWF2 opsin expression and significantly less abundant expression of other LWS opsin types in immature and mature males relative to teneral males (Fig. 7). Visual modelling, informed by *in vitro* measurements of opsin sensitivity, indicated that visual systems with maximal LWS opsin sensitivity set as the LWF2 λ_max_ (i.e., 543 nm) would likely outperform chromatic visual systems modelled with shorter LWS λ_max_ values (i.e., LWF1 and ERG λ_max_) in discriminating between thorax coloration of androchrome and *infuscans* females, and for detecting androchromes against vegetation. Additionally, males may respond to achromatic cues, with a nearly 2-fold difference in the overall brightness of androchrome and *infuscans* females at the descending midpoint limb of absorbance of LWS opsins around 600 nm (Fig. 3). These results suggest that abundantly expressed LWS opsins sensitive to longer wavelengths, can provide an ecological advantage for specific visual tasks involved in conspecific detection and likely discrimination between brightness and/or chromatic cues.

For mature males across all sites, the LWF2 opsin expression pattern exhibited a strong positive quadratic relationship (α>1), meaning that LWF2 expression was significantly greater in populations with either very low (<0.6) or very high (>0.8) female androchrome frequencies (Fig. 8). We established that the long wavelength shifted spectral sensitivity of LWF2 can, in principle, improve chromatic discrimination of androchrome and *infuscans* females and improve detection of androchrome females against vegetation, which could benefit males in populations were androchrome females are more or less abundant compared to other sites (although still the majority morph in all populations studied here). Further behavioural experiments will be important to causally establish if and how female morph frequencies might impact male behavioural choice.

Links between opsin expression and behaviour, while weak in some cases (Fuller et al., 2010; Wright et al., 2020) have been established in several biological systems (Sakai et al., 2018; Seehausen et al., 2008). Such studies include female guppies from low-predator populations which have been shown to prefer males with more orange/red colouration, a preference linked to higher expression of opsins sensitive to orange and red wavelengths compared to females from high-predator populations (Sandkam et al., 2015). Future work directly assessing the effect of mating outcomes on specific opsin expression profiles will provide valuable insights into the function of opsin plasticity in mate detection, mate choice and female morph preference in Odonata.

## CONCLUSION

Our study has revealed that the visual system of *I. elegans* is characterized by multiple SWS and LWS opsin genes, and that their absorbance spectra confer broad spectral sensitivity. Future studies should further assess regional photoreceptor spectral sensitivity, particularly in the ventral eye, which will reveal the level of functional redundancy and chromacity. We found striking variation in opsin gene expression profiles between male maturation stages and across populations with varying female morph frequencies. Specifically, the long-wavelength sensitive LWF2 opsin, showed expression profiles that consistently increased over male ontogeny, and showed expression changes that correlated with local female morph frequencies encountered by mate-searching mature male *I. elegans*. Conversely, the relative and proportional expression of another long-wavelength sensitive opsin, LWF1, decreased over male ontogeny. Functional analyses revealed that the range of SWS and LWS spectral sensitivity of *I. elegans* likely allows males to discriminate androchrome and *infuscans* female body reflectance, with visual modeling indicating that long-wavelength sensitivity characterized by LWF2 would improve detection of andromorphs against vegetation and discrimination between the two most common morph types. Other behavioural and physiological differences between androchrome and gynochrome females, such as aggression or fecundity differences, or ecological factors such as local lightning environments that are not necessarily related to mating behaviours may also play a role in modulating male opsin gene expression profiles. Overall, our new results suggest that directional changes in opsin expression may reflect visual plasticity that might aid males in mate recognition, morph detection and/or discrimination. Future work should aim to complement these findings from natural populations with experimental manipulations to determine if there is a casual role in mate choice in modulating expression profiles.

