## Supplemental Methods for "Opsin gene expression plasticity and spectral sensitivity in male damselflies could mediate female colour morph detection"

Opsin heterologous expression

We retrieved visual opsin open reading frame (ORF) sequences for *I. elegans* (Supplementary File S1) by mining whole genome data from the genome assembly iolscEleg1.1., using opsin sequences characterized from *Ischnura asiatica* (Futahashi et al., 2015) as a reference set. Opsin transcript and protein sequences were predicted with AUGUSTUS (Keller et al., 2011) followed by sequence annotation in HMMER v3.1b2 (<http://hmmer.org>). (Futahashi et al., 2015). Opsin ORFs flanked with suitable restriction sites for subcloning in the pcDNA5-FLAG-T2A-mruby2 expression cassette were synthesized for human codon optimization, overexpressed by transient transfection following the PaSHE procedure (Liénard et al., 2021, 2022) followed by purification and ultraviolet-visible spectroscopy (see supplemental methods). We did not include LWA1 or LWC1 since these opsin types are primarily expressed in larval tissues in other surveyed Odonates (Futahashi et al., 2015).

We first performed small scale transfections to verify that each final pcDNA5-Opsin construct produced detectable protein levels (Fig. S1). For this, HEK293T cells were seeded at a density of 0.6 x 10^6^ cells in DMEM medium (Gibco) in 6-well plates. The next day cells were transfected with a mixture of 4 ug plasmid DNA and a 1:3 ratio of PEI (1mg/mL) in Opti-Mem (Gibco). Forty-eight hours later, cells were visualized under a fluorescent microscope to verify expression of mRuby2 and then harvested in cold D-PBS (Sigma-Aldrich). Cells were resuspended in cold Ripa buffer supplemented with 1% n-Dodecyl-ß-D-maltoside and complete ethylenediaminetetraacetic acid (EDTA)-free protein inhibitors (Sigma-Aldrich), and incubated at 4°C for 1 hour, followed by western blot analysis.

For large scale opsin purifications, fifteen HEK293T cell dishes containing 2 x 10^6^ cells were transfected with 24 ug DNA and 72 uL PEI each. Five micromolar 11,*cis*-retinal was added 6hr post-transfection with fresh culture medium. Cells were collected under dim red light 48h post-transfection and decanted in cold Hepes wash buffer (3 mM MgCl2, 140 mM NaCl, 50 mM Hepes, EDTA-free protein inhibitors) followed by incubation at 4°C for 1 hour in presence of 40 µM 11,*cis*-retinal. Cell pellets were collected by ultracentrifugation and membrane proteins were solubilized for 1hr at 4°C in ice-cold extraction buffer (3 mM MgCl2, 140 mM NaCl, 50 mM Hepes, 20% glycerol vol/vol, 1% n-dodecyl β-D- maltoside, complete EDTA-free protein inhibitors) and ultracentrifuged. The crude extract (supernatant) was incubated in the dark with FLAG-resin on a rotator at 10 rpm until the next day. The crude extract containing resin-bound FLAG-epitope rhodopsin complexes was loaded onto a Pierce centrifuge purification column, washed with 3x 3 mL wash buffer (3 mM MgCl2, 140 mM NaCl, 50 mM Hepes, 20% glycerol vol/vol, 0.1% n-dodecyl β-D- maltoside) and eluted using FLAG-peptide. The eluate was concentrated to 100-120 microliters using an Amicon Ultra-2 10 kDa Amicon centrifugal filter for 50 min at 4°C and UV-visible absorption spectra (200 to 800 nm) of dark-adapted purified proteins were measured in the dark from 1.5-μL aliquots using a NanoDrop 2000/2000c UV-VIS spectrophotometer (Thermo Fisher). For each rhodopsin, we obtained estimates of lambda max through nonlinear least-square fitting to the absorbance data according to a visual template (Govardovskii et al., 2000) and performed 1000 bootstrap replication to calculate lambda max predictions and confidence intervals (Liénard et al., 2022) in R v.3.6.6.(*R Core Team*, 2023) using the packages rsample and tidymodels (Frick et al., 2022; Kuhn & Wickham, 2020).

Homology modelling

The Iel_SWb1 and Iel_LWF1 opsin amino acid sequences were uploaded to the SWISS-MODEL protein recognition engine (Waterhouse et al., 2018) to generate a template aligned against the invertebrate jumping spider rhodopsin crystal structure (PDB 6i9k) (Varma et al., 2019). The predicted homology model for each opsin was visualized and analysed in Pymol (Schrödinger, LLC, 2015) to identify sites in the *11*-cis-retinal binding pocket within a range of 5Å from any carbon in the retinal polyene chain.

Quantitative PCR analysis of *I. elegans* opsin expression levels

RNA was extracted from the heads of male *I. elegans* using a Monarch Total RNA Miniprep Kit protocol (Thermo Fisher Scientific, Waltham, MA, USA) following the manufacturer’s instructions and including a genomic DNA clean-up. cDNA was synthesized for each sample using a GoScript™ Reverse Transcriptase kit (Promega, Madison, WI, USA). The *Ischnura*-specific RL14 and GADPH gene sequences were annotated as described above for opsin sequences and used as reference for mRNA quantification.

For quantitative PCR analysis, two sets of gene-specific primers were designed in Geneious Prime v.2022.91 (Biomatters, Auckland, New Zealand) for both housekeeping genes and UV, SWb1, SWb2, LWA2, LWF1 – F4, and LWE1 opsins. We did not include LWA1 or LWC1 since these opsin types are primarily expressed in larval tissues (Futahashi et al., 2015). We tested the efficiency of designed primers by generating a standard curve following a 4-fold dilution of *I. elegans* cDNA and selected one primer pair for each opsin based on amplification efficiency (Table S10). The threshold detection cycle (Ct) values of selected primers were plotted against the log of cDNA dilutions and the efficiency percentage values were assessed to ensure that they were within acceptable ranges (94 – 117%) and R^2^ (0.95 – 1.00). PCR amplicons were run by electrophoresis on a 2% agarose gel to ensure the absence of primer-dimers and PCR samples for each primer pair-opsin target were purified using Exonuclease (EXO) and Shrimp alkaline phosphatase (SAP) (NEB) prior to Sanger Sequencing to confirm gene-specific amplification. Hereafter, we use abbreviated names for the opsins SWb1, SWb2, LWA2, and LWE1, which have been shortened to SW1, SW2, LWA, and LWE, respectively.

qPCR was performed in a CFX384 Real-Time System (BioRad Laboratories, Inc., Hercules, CA, USA). Reaction mixes were prepared that contained 1x SYBR Green, 1uM primer mix, and 0.1 ng cDNA. The program was designed as follows: denaturation at 95°C for 2min, followed by 39 cycles at 95°C for 10s and 60°C for 10s, with melting curve analysis performed from 65.5°C to 89.5°C at 0.6°C increments to confirm the specificity of PCR products. Two independent qPCR assays (technical assays) were processed for each combination of a given cDNA sample and opsin-specific primer pair, and triplicate wells were run within each qPCR assay. We averaged Ct values within each assay for each opsin target (average Ct values = 2 per {male sample x opsin gene target}.

Visual modelling of female morph detection and discrimination

To test whether expression of different LWS opsins might impact detection of females against the background or discrimination between andromorph and infuscans females, we performed visual modelling using custom MATLAB scripts (ver. 9.9.0.1592791 (R2020b) Update 5). We calculated just noticeable differences (JNDs) using a receptor noise limited model (Vorobyev & Osorio, 1998), which allows robust comparisons on JNDs between several parsimonious visual systems, with JND values of 1 indicating the threshold for discrimination between two stimuli.

We calculated JNDs for three possible visual Odonate systems with varying maximal sensitivities for the SWS and LWS opsins, whereas the UVS λ_max_ was set to 360 nm in all models, based on electroretinograms (ERG) in *Sympetrum frequens* (Futahashi et al 2015) and *Ischnura* *heterosticta* (Huang et al. 2014). The first modelled visual system follows ERG spectral sensitivity values from *Ischnura heterosticta* with SWS λ_max_ = 450 nm and LWS λ_max_ = 525 nm (Huang et al., 2014). For the second and third visual models, we calculated JNDs with SWS λ_max_ = 413 nm, mean value of *I. elegans* SWb1 and SWb2 opsin sensitivities measured *in vitro*, together with LWS λ_max_ = 531 nm (model 2) or 543 nm (model 3), corresponding to maximal in vitro absorbance of LWF1 and LWF2 opsins, respectively. We focused on LWF1 and LWF2 opsins owing to significant decreased expression of LWF1 and increased expression of LWF2 in mature males relative to immature males (Fig. 8). Spectral sensitivity curves for UVS, SWS, and LWS receptors were generated following a Govardovskii template (Govardovskii et al., 2000).

We first calculated receptor quantum catch, *q*, for each stimulus using the following equation:

$$q_{i}=\int_{300}^{700} R_{i}\left( \lambda\right)S\left( \lambda\right)d\lambda$$

where *R* is the sensitivity of receptor *i*, and *S* is stimulus reflectance. We used reflectance data of mature female thorax coloration for androchrome and infuscans from (Henze et al., 2019) and spectra from green and brown vegetation from (Henze et al., 2018) as stimuli.

Next, the contrast for each receptor channel is calculated as:

$${\Delta f}_{i}=ln\frac{q_{i_{stim1}}}{q_{i_{stim2}}}$$

Estimates of receptor noise in bright light conditions are equivalent to the Weber fraction (*⍵*), scaled by the relative abundance of receptor types in the eye. We calculated noise as:

$$e_{i}=\sqrt{\frac{nL}{\eta_{i}}eL}$$

where *e* is noise in the receptor channel, and η is the relative density of a particular receptor type in the retina. We used a noise value of 0.1 for the LWS channel and a receptors ratio of 1 UVS: 1 SWS: 7 LWS.

Finally chromatic contrasts (JNDs) are calculated for a trichromatic visual system as:

$$\Delta S^{2}=\frac{{e_{1}}^{2}\left( {\Delta f}_{3}-{\Delta f}_{2} \right)^{2}+{e_{2}}^{2}\left( {\Delta f}_{3}-{\Delta f}_{1} \right)^{2}+{e_{3}}^{2}{({\Delta f}_{2}-{\Delta f}_{1})}^{2}}{\left( e_{1}e_{2} \right)^{2}+\left( e_{1}e_{3} \right)^{2}+\left( e_{2}e_{3} \right)^{2}}$$

Statistical analysis

Model selection was performed using the lme4 package (Bates et al., 2015) in R v 4.1.1. For model selection, we first selected the random effect structure by comparing Akaike information criterions (AICs) between models fitted using restricted maximum likelihood (REML) estimation that included all possible fixed effects and relevant interactions. Following selection of the random effect structure, we used backward model selection with maximum likelihood (ML) estimation to determine the optimal fixed effect structure by testing the significance of each interaction or term. After selection of the optimal random and fixed effect structures, the final model was run using REML estimation and the model output and results of post-hoc analysis were obtained using the lmerTest and emmeans packages (Kuznetsova et al., 2017; Lenth, 2022), respectively.

*Opsin mRNA expression in teneral, immature, and mature males across sites*

We tested the effect of maturity stage (factor with 3 levels: teneral, immature, mature), opsin type (factor with 8 levels: SW1, SW2, LWA, LWF1 – LWF4, LWE), and their interaction on log relative opsin expression. Model selection indicated a significant interaction between maturity level and opsin type (likelihood ratio test: χ2 (14) = 163.9, *p* < 0.0001). The final model therefore included year, qPCR replicate, and male ID as random effects with fixed effects of maturity stage, opsin type, and their interaction to explain log relative opsin expression in male *I. elegans*.

        	To investigate how proportional opsin gene expression changes over development, we again used maturity stage, opsin type, and their interaction as fixed effects, but now only focused on opsins found in the compound eye (factor with 7 levels: SW1, SW2, LWA, LWF1 – LWF4) with beta regression model using the package glmmTMB (Brooks et al., 2023). Data exploration suggested violations of the homogeneity of variance assumption for this dataset. We therefore allowed for differing variance structure in the terms *maturity stage* and *opsin* using ‘dispformula’ term in the glmmTMB package. We included only male ID as a random effect since inclusion of additional random effects did not improve model AIC. Model selection showed a significant interaction between opsin type and maturity stage (likelihood ratio test: χ2 (12) = 22196, *p* < 0.0001), making the final model the interaction of maturity stage and opsin type on proportion opsin expression.

Preliminary analysis suggested differences in relative opsin expression between teneral males from the two field populations and those from outdoor mesocosm tanks at Stensoffa Field Station (Table S11). Despite these differences, the results with and without males from the mesocosm tanks were largely comparable. We therefore retained the teneral males from the mesocosm tanks in our main analyses. Results of log relative expression and proportion opsin expression without the mesocosm males are presented in Supplementary Material (Tables S11 -S14, Fig. S4).

*Opsin mRNA expression in populations with different female morph structure*

To assess the effect of local female morph frequency on male opsin expression, we used the frequency of androchrome females for each site and year to describe local population morph frequency. We used the proportion of androchrome females within a site for each year separately, instead of generating a site average, since a primary aim of the current study was to assess if opsin expression changes in response to current local conditions. Therefore, “population morph frequency” refers to the local morph frequency in a given year, with two population morph frequencies per site.

We followed the same modeling procedure to test the effect of maturity stage, opsin type, the proportion of androchrome females, and their interactions on relative opsin expression. We included the proportion of androchrome females also as a quadratic term to account for any non-linear effect of androchrome frequencies. In this analysis, we focused only on immature and mature males since data for teneral males was not available for most sites. Backward model selection of fixed effect structure indicated that the three-way interaction between opsin type, maturity stage, and androchrome proportion was significant (likelihood ratio test: χ2 (14) = 43.9, *p* < 0.0001). The final model therefore accounted for a quadratic relationship between androchrome proportion, opsin type, maturity stage, and their interaction as fixed effects with year, male ID, and replicate as random effects.

*Opsin expression in response to mating outcomes*

We recorded the female morph of all males collected in copula in 2022 to assess whether relative opsin expression differed between males collected in copula with androchrome or infuscans females. The selected random effect structure included male ID and replicate. We considered maturity level, opsin type, and female morph (factor with 2 levels: androchrome, infuscans), and their interactions as fixed effects. Backwards model selection resulted in a final model with opsin as the only fixed effect, suggesting that there is no significant effect of female morph type on a males’ relative opsin expression in the current study. However, it is possible this is due to relatively small sample size, with n = 8 males collected in copula with an androchrome female and n = 7 males collected in copula with an infuscans female.
