## Supplemental Tables&Figures for "Opsin gene expression plasticity and spectral sensitivity in male damselflies could mediate female colour morph detection"

**Table S1.** Total counts of immature and mature female *I. elegans* collected across sites and years.

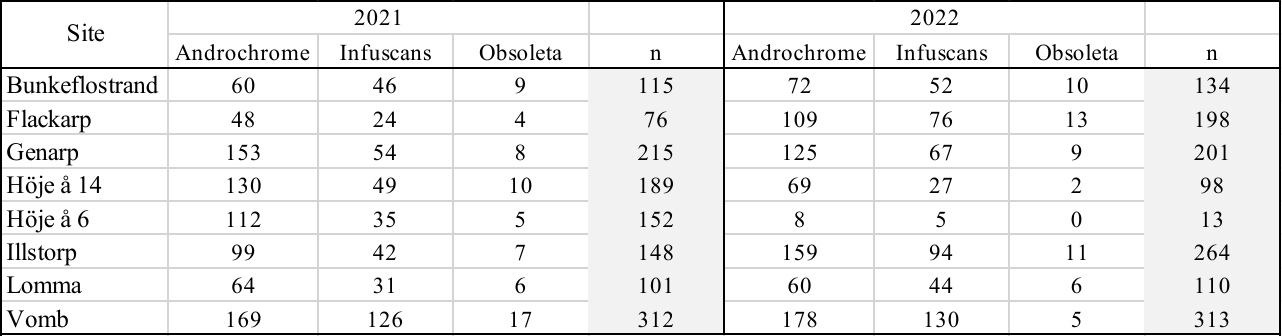

**Table S2**. Number of collected male *I. elegans* that were included in qPCR data analysis from eight natural sites and semi-naturalistic mesocosm tanks at Stensoffa Field Station across two collection years. The number in parentheses shows the number of males that were paired with a female (*in copula*) when collected.

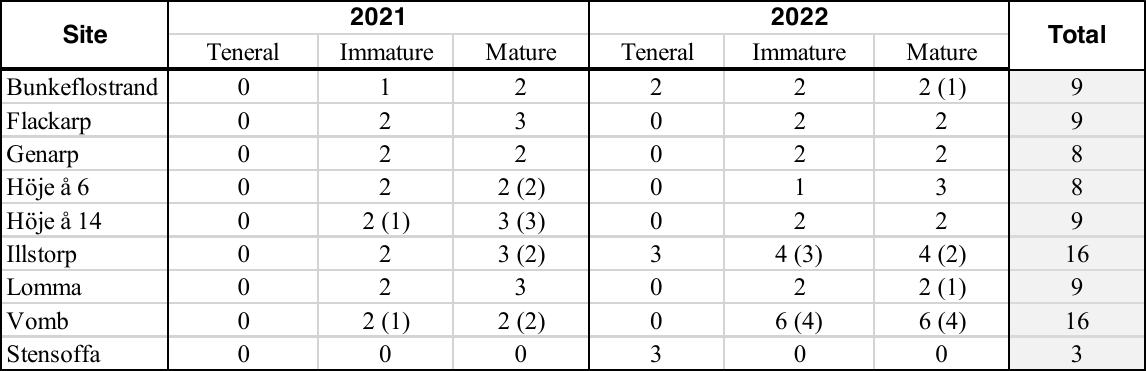

**Table S3.** Number of females expressing mature body coloration for each site across years and results of a Fisher’s Exact Test for differences in female morph frequency across years.

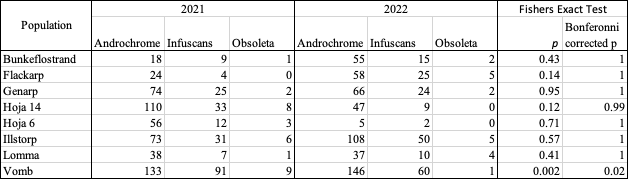

**Table S4.** *I. elegans* opsin amino acid sites predicted to lie within 5A of the cis-retinal chromophore binding pocket.

| **Residue location** | **SWB1** | | **SWB2** | | |
| --- | --- | --- | --- | --- | --- |
|  | **Amino acid position*** | **Amino acid** | | **Amino acid position*** | **Amino acid** |
| TM2 | 112 | S | | 118 | S |
| **TM3** | **136** | **Y** | | **142** | **F** |
| TM3 | 137 | G | | 143 | G |
| TM3 | 140 | G | | 146 | G |
| TM3 | 141 | S | | 147 | S |
| TM3 | 144 | G | | 150 | G |
| TM3 | 145 | M | | 151 | M |
| beta-sheet | 201 | Y | | 207 | Y |
| beta-sheet | 204 | E | | 210 | E |
| beta-sheet | 209 | T | | 215 | T |
| beta-sheet | 210 | C | | 216 | C |
| beta-sheet | 211 | S | | 217 | S |
| beta-sheet | 212 | F | | 218 | F |
| TM5 | 228 | I | | 234 | I |
| TM5 | 229 | F | | 235 | F |
| TM5 | 233 | Y | | 239 | Y |
| **TM6** | **294** | **Y** | | **300** | **F** |
| TM6 | 298 | W | | 304 | W |
| TM7 | 301 | Y | | 307 | Y |
| TM8 | 302 | A | | 308 | A |
| TM6 | 305 | A | | 311 | A |
| TM7 | 325 | A | | 331 | A |
| TM7 | 335 | K | | 335 | K |
| **Residue location** | **LWF1-LWF4** | | **LWA2** | | |
|  | **Amino acid position**** | **Amino acid** | | **Amino acid position** Amino acid** | |
| TM2 | 107 | M | | 107 | M |
| TM3 | 130 | Y | | 130 | Y |
| **TM3** | **131** | **A** | | **131** | **G** |
| TM3 | 134 | G | | 134 | G |
| TM3 | 135 | S | | 135 | S |
| TM3 | 138 | G | | 138 | G |
| TM3 | 139 | C | | 139 | C |
| beta-sheet | 195 | Y | | 195 | Y |
| beta-sheet | 198 | E | | 198 | E |
| beta-sheet | 204 | C | | 204 | C |
| beta-sheet | 205 | G | | 205 | G |
| beta-sheet | 206 | T | | 206 | T |
| TM5 | 222 | Y | | 222 | Y |
| TM5 | 223 | S | | 223 | S |
| TM5 | 226 | C | | 226 | C |
| TM5 | 227 | Y | | 227 | Y |
| TM6 | 289 | W | | 289 | W |
| TM6 | 293 | W | | 293 | W |
| TM6 | 296 | Y | | 296 | Y |
| TM6 | 297 | L | | 297 | L |
| TM6 | 300 | N | | 300 | N |
| **TM7** | **319** | **S** | | **319** | **A** |
| TM7 | 322 | A | | 322 | A |
| TM7 | 323 | K | | 323 | K |

* based on original position in unaligned sequence

** based on position in multiple alignment with LWF1-F4 and LWA

**Table S5.** Post-hoc test with Tukey correction for relative opsin expression within each maturity level tested. *P*-values significant at an alpha level of 0.05 or below are shown in bold.

| Teneral | | | | | |
| --- | --- | --- | --- | --- | --- |
| contrast | estimate | SE | df | t.ratio | p.value |
| LWA - LWE | 1.2889 | 0.1656 | 1186 | 7.785 | **<.0001** |
| LWA - LWF1 | 0.1734 | 0.1656 | 1186 | 1.048 | 0.9669 |
| LWA - LWF2 | -0.0345 | 0.1656 | 1186 | -0.209 | 1 |
| LWA - LWF3 | 0.2252 | 0.1656 | 1186 | 1.36 | 0.875 |
| LWA - LWF4 | -0.605 | 0.1656 | 1186 | -3.655 | **0.0065** |
| LWA - SW1 | 0.8337 | 0.1656 | 1186 | 5.036 | **<.0001** |
| LWA - SW2 | 1.1261 | 0.1656 | 1186 | 6.802 | **<.0001** |
| LWE - LWF1 | -1.1154 | 0.1656 | 1186 | -6.737 | **<.0001** |
| LWE - LWF2 | -1.3234 | 0.1656 | 1186 | -7.994 | **<.0001** |
| LWE - LWF3 | -1.0636 | 0.1656 | 1186 | -6.425 | **<.0001** |
| LWE - LWF4 | -1.8939 | 0.1656 | 1186 | -11.44 | **<.0001** |
| LWE - SW1 | -0.4552 | 0.1656 | 1186 | -2.75 | 0.1091 |
| LWE - SW2 | -0.1628 | 0.1656 | 1186 | -0.983 | 0.9768 |
| LWF1 - LWF2 | -0.208 | 0.1656 | 1186 | -1.256 | 0.9145 |
| LWF1 - LWF3 | 0.0518 | 0.1656 | 1186 | 0.313 | 1 |
| LWF1 - LWF4 | -0.7785 | 0.1656 | 1186 | -4.702 | **0.0001** |
| LWF1 - SW1 | 0.6602 | 0.1656 | 1186 | 3.988 | **0.0018** |
| LWF1 - SW2 | 0.9526 | 0.1656 | 1186 | 5.754 | **<.0001** |
| LWF2 - LWF3 | 0.2597 | 0.1656 | 1186 | 1.569 | 0.7689 |
| LWF2 - LWF4 | -0.5705 | 0.1656 | 1186 | -3.446 | **0.0137** |
| LWF2 - SW1 | 0.8682 | 0.1656 | 1186 | 5.244 | **<.0001** |
| LWF2 - SW2 | 1.1606 | 0.1656 | 1186 | 7.01 | **<.0001** |
| LWF3 - LWF4 | -0.8302 | 0.1656 | 1186 | -5.015 | **<.0001** |
| LWF3 - SW1 | 0.6084 | 0.1656 | 1186 | 3.675 | **0.006** |
| LWF3 - SW2 | 0.9009 | 0.1656 | 1186 | 5.442 | **<.0001** |
| LWF4 - SW1 | 1.4387 | 0.1656 | 1186 | 8.69 | **<.0001** |
| LWF4 - SW2 | 1.7311 | 0.1656 | 1186 | 10.456 | **<.0001** |
| SW1 - SW2 | 0.2924 | 0.1656 | 1186 | 1.766 | 0.6432 |
| Immature | | | | | |
| contrast | estimate | SE | df | t.ratio | p.value |
| LWA - LWE | 1.7525 | 0.078 | 1186 | 22.456 | **<.0001** |
| LWA - LWF1 | 0.8491 | 0.0783 | 1187 | 10.84 | **<.0001** |
| LWA - LWF2 | -0.9342 | 0.078 | 1186 | -11.97 | **<.0001** |
| LWA - LWF3 | 0.2579 | 0.078 | 1186 | 3.304 | **0.0219** |
| LWA - LWF4 | -0.6256 | 0.078 | 1186 | -8.016 | **<.0001** |
| LWA - SW1 | 0.4482 | 0.078 | 1186 | 5.742 | **<.0001** |
| LWA - SW2 | 0.8611 | 0.078 | 1186 | 11.034 | **<.0001** |
| LWE - LWF1 | -0.9034 | 0.0783 | 1187 | -11.532 | **<.0001** |
| LWE - LWF2 | -2.6867 | 0.078 | 1186 | -34.426 | **<.0001** |
| LWE - LWF3 | -1.4946 | 0.078 | 1186 | -19.152 | **<.0001** |
| LWE - LWF4 | -2.3781 | 0.078 | 1186 | -30.472 | **<.0001** |
| LWE - SW1 | -1.3044 | 0.078 | 1186 | -16.713 | **<.0001** |
| LWE - SW2 | -0.8914 | 0.078 | 1186 | -11.422 | **<.0001** |
| LWF1 - LWF2 | -1.7833 | 0.0783 | 1187 | -22.765 | **<.0001** |
| LWF1 - LWF3 | -0.5913 | 0.0783 | 1187 | -7.548 | **<.0001** |
| LWF1 - LWF4 | -1.4747 | 0.0783 | 1187 | -18.826 | **<.0001** |
| LWF1 - SW1 | -0.401 | 0.0783 | 1187 | -5.119 | **<.0001** |
| LWF1 - SW2 | 0.012 | 0.0783 | 1187 | 0.153 | 1 |
| LWF2 - LWF3 | 1.192 | 0.078 | 1186 | 15.274 | **<.0001** |
| LWF2 - LWF4 | 0.3086 | 0.078 | 1186 | 3.954 | **0.0021** |
| LWF2 - SW1 | 1.3823 | 0.078 | 1186 | 17.712 | **<.0001** |
| LWF2 - SW2 | 1.7953 | 0.078 | 1186 | 23.004 | **<.0001** |
| LWF3 - LWF4 | -0.8834 | 0.078 | 1186 | -11.32 | **<.0001** |
| LWF3 - SW1 | 0.1903 | 0.078 | 1186 | 2.438 | 0.2238 |
| LWF3 - SW2 | 0.6033 | 0.078 | 1186 | 7.73 | **<.0001** |
| LWF4 - SW1 | 1.0737 | 0.078 | 1186 | 13.758 | **<.0001** |
| LWF4 - SW2 | 1.4867 | 0.078 | 1186 | 19.05 | **<.0001** |
| SW1 - SW2 | 0.413 | 0.078 | 1186 | 5.292 | **<.0001** |
| Mature | | | | | |
| contrast | estimate | SE | df | t.ratio | p.value |
| LWA - LWE | 1.8662 | 0.0729 | 1190 | 25.608 | **<.0001** |
| LWA - LWF1 | 1.111 | 0.0724 | 1189 | 15.353 | **<.0001** |
| LWA - LWF2 | -1.1679 | 0.0714 | 1186 | -16.355 | **<.0001** |
| LWA - LWF3 | 0.2273 | 0.0714 | 1186 | 3.183 | **0.0322** |
| LWA - LWF4 | -0.657 | 0.0714 | 1186 | -9.201 | **<.0001** |
| LWA - SW1 | 0.3706 | 0.0714 | 1186 | 5.19 | **<.0001** |
| LWA - SW2 | 0.8675 | 0.0714 | 1186 | 12.148 | **<.0001** |
| LWE - LWF1 | -0.7552 | 0.0736 | 1188 | -10.259 | **<.0001** |
| LWE - LWF2 | -3.034 | 0.0729 | 1190 | -41.633 | **<.0001** |
| LWE - LWF3 | -1.6389 | 0.0729 | 1190 | -22.489 | **<.0001** |
| LWE - LWF4 | -2.5232 | 0.0729 | 1190 | -34.623 | **<.0001** |
| LWE - SW1 | -1.4956 | 0.0729 | 1190 | -20.523 | **<.0001** |
| LWE - SW2 | -0.9987 | 0.0729 | 1190 | -13.704 | **<.0001** |
| LWF1 - LWF2 | -2.2788 | 0.0724 | 1189 | -31.492 | **<.0001** |
| LWF1 - LWF3 | -0.8837 | 0.0724 | 1189 | -12.212 | **<.0001** |
| LWF1 - LWF4 | -1.768 | 0.0724 | 1189 | -24.433 | **<.0001** |
| LWF1 - SW1 | -0.7404 | 0.0724 | 1189 | -10.232 | **<.0001** |
| LWF1 - SW2 | -0.2435 | 0.0724 | 1189 | -3.365 | **0.0179** |
| LWF2 - LWF3 | 1.3951 | 0.0714 | 1186 | 19.538 | **<.0001** |
| LWF2 - LWF4 | 0.5109 | 0.0714 | 1186 | 7.154 | **<.0001** |
| LWF2 - SW1 | 1.5384 | 0.0714 | 1186 | 21.544 | **<.0001** |
| LWF2 - SW2 | 2.0353 | 0.0714 | 1186 | 28.503 | **<.0001** |
| LWF3 - LWF4 | -0.8843 | 0.0714 | 1186 | -12.384 | **<.0001** |
| LWF3 - SW1 | 0.1433 | 0.0714 | 1186 | 2.007 | 0.4776 |
| LWF3 - SW2 | 0.6402 | 0.0714 | 1186 | 8.965 | **<.0001** |
| LWF4 - SW1 | 1.0276 | 0.0714 | 1186 | 14.39 | **<.0001** |
| LWF4 - SW2 | 1.5245 | 0.0714 | 1186 | 21.349 | **<.0001** |
| SW1 - SW2 | 0.4969 | 0.0714 | 1186 | 6.958 | **<.0001** |

**Table S6.** Post-hoc analysis for relative opsin expression with Tukey correction between teneral (T), immature (I), and mature (M) male *I. elegans* for each opsin class. *P*-values significant at an alpha level of 0.05 of below are shown in bold.

| Opsin = SW1: |  |  |  |  |  |
| --- | --- | --- | --- | --- | --- |
| contrast | estimate | SE | df | t.ratio | p.value |
| I - M | -0.1156 | 0.0916 | 419 | -1.262 | 0.4176 |
| I - T | 0.3064 | 0.1606 | 401 | 1.908 | 0.1378 |
| M - T | 0.422 | 0.1588 | 396 | 2.657 | **0.0223** |
| Opsin = SW2: | | | | | |
| contrast | estimate | SE | df | t.ratio | p.value |
| I - M | -0.0317 | 0.0916 | 419 | -0.346 | 0.9361 |
| I - T | 0.1858 | 0.1606 | 401 | 1.157 | 0.4797 |
| M - T | 0.2175 | 0.1588 | 396 | 1.37 | 0.3579 |
| Opsin = LWA: | | | | | |
| contrast | estimate | SE | df | t.ratio | p.value |
| I - M | -0.038 | 0.0916 | 419 | -0.415 | 0.9093 |
| I - T | -0.0791 | 0.1606 | 401 | -0.493 | 0.8749 |
| M - T | -0.0411 | 0.1588 | 396 | -0.259 | 0.9638 |
| Opsin = LWF1: | | | | | |
| contrast | estimate | SE | df | t.ratio | p.value |
| I - M | 0.2238 | 0.0926 | 433 | 2.417 | **0.0424** |
| I - T | -0.7548 | 0.1607 | 402 | -4.696 | **<.0001** |
| M - T | -0.9786 | 0.1592 | 399 | -6.148 | **<.0001** |
| Opsin = LWF2: | | | | | |
| contrast | estimate | SE | df | t.ratio | p.value |
| I - M | -0.2717 | 0.0916 | 419 | -2.966 | **0.0089** |
| I - T | 0.8205 | 0.1606 | 401 | 5.109 | **<.0001** |
| M - T | 1.0922 | 0.1588 | 396 | 6.877 | **<.0001** |
| Opsin = LWF3: | | | | | |
| contrast | estimate | SE | df | t.ratio | p.value |
| I - M | -0.0686 | 0.0916 | 419 | -0.749 | 0.7344 |
| I - T | -0.1118 | 0.1606 | 401 | -0.696 | 0.766 |
| M - T | -0.0432 | 0.1588 | 396 | -0.272 | 0.9601 |
| Opsin = LWF4: | | | | | |
| contrast | estimate | SE | df | t.ratio | p.value |
| I - M | -0.0695 | 0.0916 | 419 | -0.758 | 0.7288 |
| I - T | -0.0586 | 0.1606 | 401 | -0.365 | 0.9293 |
| M - T | 0.0109 | 0.1588 | 396 | 0.069 | 0.9974 |
| Opsin = LWE: | | | | | |
| contrast | estimate | SE | df | t.ratio | p.value |
| I - M | 0.0756 | 0.0927 | 435 | 0.815 | 0.6937 |
| I - T | -0.5428 | 0.1606 | 401 | -3.38 | **0.0023** |
| M - T | -0.6184 | 0.1594 | 400 | -3.88 | **0.0004** |

**Table S7**. Post-hoc analysis with Tukey correction for relative opsin expression over adult development for male *I. elegans* by site. Significant differences are shown in bold. No significant interaction between maturity stage and opsin type was indicated for Höje å 14 (df = 7, F = 1.2, *p* = 0.32), Vomb (df = 7, F = 1.6, *p* = 0.14), and Lomma (df = 7, F = 0.66, *p* = 0.70), and so post-hoc comparisons for these sites are not included below.

| Bunkeflostrand | | | | | |
| --- | --- | --- | --- | --- | --- |
| **SW1** | estimate | SE | df | t.ratio | p.value |
| Teneral - Immature | -0.2414 | 0.145 | 98.6 | -1.665 | 0.224 |
| Teneral -Mature | -0.3627 | 0.139 | 94.1 | -2.616 | **0.0277** |
| Immature - Mature | -0.1213 | 0.121 | 100.5 | -1.003 | 0.5764 |
| **SW2** | estimate | SE | df | t.ratio | p.value |
| Teneral - Immature | -0.0212 | 0.145 | 98.6 | -0.147 | 0.9882 |
| Teneral -Mature | -0.1303 | 0.139 | 94.1 | -0.94 | 0.6167 |
| Immature - Mature | -0.1091 | 0.121 | 100.5 | -0.902 | 0.6403 |
| **LWA** | estimate | SE | df | t.ratio | p.value |
| Teneral - Immature | 0.2747 | 0.145 | 98.6 | 1.895 | 0.1456 |
| Teneral -Mature | 0.0318 | 0.139 | 94.1 | 0.229 | 0.9714 |
| Immature - Mature | -0.2429 | 0.121 | 100.5 | -2.009 | 0.1152 |
| **LWF1** | estimate | SE | df | t.ratio | p.value |
| Teneral - Immature | 0.9488 | 0.145 | 98.6 | 6.543 | **<.0001** |
| Teneral -Mature | 1.3687 | 0.139 | 94.1 | 9.871 | **<.0001** |
| Immature - Mature | 0.4199 | 0.121 | 100.5 | 3.473 | **0.0022** |
| **LWF2** | estimate | SE | df | t.ratio | p.value |
| Teneral - Immature | -0.4898 | 0.145 | 98.6 | -3.377 | **0.003** |
| Teneral -Mature | -0.8101 | 0.139 | 94.1 | -5.842 | **<.0001** |
| Immature - Mature | -0.3203 | 0.121 | 100.5 | -2.649 | **0.0252** |
| **LWF3** | estimate | SE | df | t.ratio | p.value |
| Teneral - Immature | 0.2956 | 0.145 | 98.6 | 2.038 | 0.1084 |
| Teneral -Mature | 0.0143 | 0.139 | 94.1 | 0.103 | 0.9941 |
| Immature - Mature | -0.2813 | 0.121 | 100.5 | -2.326 | 0.0567 |
| **LWF4** | estimate | SE | df | t.ratio | p.value |
| Teneral - Immature | 0.3502 | 0.145 | 98.6 | 2.415 | **0.0459** |
| Teneral -Mature | 0.1592 | 0.139 | 94.1 | 1.148 | 0.487 |
| Immature - Mature | -0.191 | 0.121 | 100.5 | -1.579 | 0.2592 |
| **LWE** | estimate | SE | df | t.ratio | p.value |
| Teneral - Immature | 0.0713 | 0.145 | 98.6 | 0.492 | 0.8755 |
| Teneral -Mature | 0.5953 | 0.139 | 94.1 | 4.293 | **0.0001** |
| Immature - Mature | 0.524 | 0.121 | 100.5 | 4.334 | **0.0001** |

| Illstorp | | | | | |
| --- | --- | --- | --- | --- | --- |
| **SW1** | estimate | SE | df | t.ratio | p.value |
| Teneral - Immature | -0.236 | 0.239 | 81.9 | -0.989 | 0.5858 |
| Teneral -Mature | -0.2566 | 0.233 | 81.9 | -1.102 | 0.5155 |
| Immature - Mature | -0.0206 | 0.188 | 81.9 | -0.11 | 0.9934 |
| **SW2** | estimate | SE | df | t.ratio | p.value |
| Teneral - Immature | 0.0956 | 0.239 | 81.9 | 0.401 | 0.9154 |
| Teneral -Mature | 0.0184 | 0.233 | 81.9 | 0.079 | 0.9966 |
| Immature - Mature | -0.0772 | 0.188 | 81.9 | -0.411 | 0.9111 |
| **LWA** | estimate | SE | df | t.ratio | p.value |
| Teneral - Immature | 0.3172 | 0.239 | 81.9 | 1.33 | 0.383 |
| Teneral -Mature | 0.3324 | 0.233 | 81.9 | 1.428 | 0.3314 |
| Immature - Mature | 0.0153 | 0.188 | 81.9 | 0.081 | 0.9964 |
| **LWF1** | estimate | SE | df | t.ratio | p.value |
| Teneral - Immature | 0.8827 | 0.239 | 81.9 | 3.7 | **0.0011** |
| Teneral -Mature | 0.7881 | 0.233 | 81.9 | 3.385 | **0.0031** |
| Immature - Mature | -0.0946 | 0.188 | 81.9 | -0.504 | 0.8697 |
| **LWF2** | estimate | SE | df | t.ratio | p.value |
| Teneral - Immature | -0.2415 | 0.239 | 81.9 | -1.012 | 0.5713 |
| Teneral -Mature | -0.568 | 0.233 | 81.9 | -2.44 | **0.044** |
| Immature - Mature | -0.3265 | 0.188 | 81.9 | -1.739 | 0.1969 |
| **LWF3** | estimate | SE | df | t.ratio | p.value |
| Teneral - Immature | 0.3422 | 0.239 | 81.9 | 1.434 | 0.3282 |
| Teneral -Mature | 0.2375 | 0.233 | 81.9 | 1.02 | 0.5664 |
| Immature - Mature | -0.1047 | 0.188 | 81.9 | -0.558 | 0.8428 |
| **LWF4** | estimate | SE | df | t.ratio | p.value |
| Teneral - Immature | 0.2682 | 0.239 | 81.9 | 1.124 | 0.5019 |
| Teneral -Mature | 0.1661 | 0.233 | 81.9 | 0.714 | 0.7561 |
| Immature - Mature | -0.1021 | 0.188 | 81.9 | -0.544 | 0.8499 |
| **LWE** | estimate | SE | df | t.ratio | p.value |
| Teneral - Immature | -0.0967 | 0.239 | 81.9 | -0.405 | 0.9136 |
| Teneral -Mature | 0.8534 | 0.233 | 81.9 | 3.666 | **0.0013** |
| Immature - Mature | 0.9501 | 0.188 | 81.9 | 5.062 | **<.0001** |

| Flackarp | | | | | |
| --- | --- | --- | --- | --- | --- |
| **SW1** | estimate | SE | df | t.ratio | p.value |
| Immature - Mature | -0.0837 | 0.309 | 26.6 | -0.271 | 0.7883 |
| **SW2** | estimate | SE | df | t.ratio | p.value |
| Immature - Mature | -0.2312 | 0.309 | 26.6 | -0.749 | 0.4602 |
| **LWA** | estimate | SE | df | t.ratio | p.value |
| Immature - Mature | 0.0813 | 0.309 | 26.6 | 0.263 | 0.7943 |
| **LWF1** | estimate | SE | df | t.ratio | p.value |
| Immature - Mature | 0.3584 | 0.313 | 28 | 1.145 | 0.262 |
| **LWF2** | estimate | SE | df | t.ratio | p.value |
| Immature - Mature | -0.794 | 0.309 | 26.6 | -2.573 | **0.016** |
| **LWF3** | estimate | SE | df | t.ratio | p.value |
| Immature - Mature | 0.2962 | 0.309 | 26.6 | 0.96 | 0.3456 |
| **LWF4** | estimate | SE | df | t.ratio | p.value |
| Immature - Mature | 0.3399 | 0.309 | 26.6 | 1.102 | 0.2804 |
| **LWE** | estimate | SE | df | t.ratio | p.value |
| Immature - Mature | -0.7576 | 0.319 | 29.8 | -2.377 | **0.0241** |

| Genarp | | | | | |
| --- | --- | --- | --- | --- | --- |
| **SW1** | estimate | SE | df | t.ratio | p.value |
| Immature - Mature | -0.06 | 0.269 | 35 | -0.205 | 0.8389 |
| **SW2** | estimate | SE | df | t.ratio | p.value |
| Immature - Mature | 0.044 | 0.269 | 35 | 0.164 | 0.8708 |
| **LWA** | estimate | SE | df | t.ratio | p.value |
| Immature - Mature | -0.2288 | 0.269 | 35 | -0.852 | 0.4002 |
| **LWF1** | estimate | SE | df | t.ratio | p.value |
| Immature - Mature | -0.2129 | 0.276 | 37.9 | -0.771 | 0.4453 |
| **LWF2** | estimate | SE | df | t.ratio | p.value |
| Immature - Mature | 0.2243 | 0.269 | 35 | 0.835 | 0.4094 |
| **LWF3** | estimate | SE | df | t.ratio | p.value |
| Immature - Mature | -0.0852 | 0.269 | 35 | -0.317 | 0.7529 |
| **LWF4** | estimate | SE | df | t.ratio | p.value |
| Immature - Mature | -0.31 | 0.269 | 35 | -1.143 | 0.2609 |
| **LWE** | estimate | SE | df | t.ratio | p.value |
| Immature - Mature | 0.9202 | 0.269 | 35 | 3.425 | **0.0016** |

| Höje å 6 | | | | | |
| --- | --- | --- | --- | --- | --- |
| **SW1** | estimate | SE | df | t.ratio | p.value |
| Immature - Mature | -0.1065 | 0.211 | 12.5 | -0.504 | 0.6232 |
| **SW2** | estimate | SE | df | t.ratio | p.value |
| Immature - Mature | -0.0494 | 0.211 | 12.5 | -0.234 | 0.8188 |
| **LWA** | estimate | SE | df | t.ratio | p.value |
| Immature - Mature | -0.2848 | 0.211 | 12.5 | -1.348 | 0.2015 |
| **LWF1** | estimate | SE | df | t.ratio | p.value |
| Immature - Mature | -0.0303 | 0.211 | 12.5 | -0.143 | 0.8883 |
| **LWF2** | estimate | SE | df | t.ratio | p.value |
| Immature - Mature | -0.0369 | 0.211 | 12.5 | -0.174 | 0.8643 |
| **LWF3** | estimate | SE | df | t.ratio | p.value |
| Immature - Mature | -0.308 | 0.211 | 12.5 | -1.457 | 0.1696 |
| **LWF4** | estimate | SE | df | t.ratio | p.value |
| Immature - Mature | -0.5373 | 0.211 | 12.5 | -2.543 | **0.0251** |
| **LWE** | estimate | SE | df | t.ratio | p.value |
| Immature - Mature | -0.3984 | 0.211 | 12.5 | -1.886 | 0.0828 |

**Table S8.** Post-hoc analysis of proportion opsin expression with Tukey correction between teneral (T), immature (I), and mature (M) male *I. elegans* for each opsin class. *P*-values significant at an alpha level of 0.05 of below are shown in bold.

| Opsin = SW1: |  |  |  |  |  |
| --- | --- | --- | --- | --- | --- |
| contrast | estimate | SE | df | t.ratio | p.value |
| I - M | 0.001884 | 0.002447 | 84 | 0.77 | 0.7224 |
| I - T | 0.001301 | 0.004234 | 84 | 0.307 | 0.9494 |
| M - T | -0.000584 | 0.004171 | 84 | -0.14 | 0.9893 |
| Opsin = SW2 |  |  |  |  |  |
| contrast | estimate | SE | df | t.ratio | p.value |
| I - M | 0.001741 | 0.000679 | 84 | 2.565 | **0.0321** |
| I - T | -0.002392 | 0.001175 | 84 | -2.036 | 0.1099 |
| M - T | -0.004134 | 0.001157 | 84 | -3.572 | **0.0017** |
| Opsin = LWA |  |  |  |  |  |
| contrast | estimate | SE | df | t.ratio | p.value |
| I - M | 0.004231 | 0.008356 | 84 | 0.506 | 0.8685 |
| I - T | -0.060963 | 0.014458 | 84 | -4.216 | **0.0002** |
| M - T | -0.065194 | 0.014243 | 84 | -4.577 | **<.0001** |
| Opsin = LWF1 |  |  |  |  |  |
| contrast | estimate | SE | df | t.ratio | p.value |
| I - M | 0.014191 | 0.003226 | 84 | 4.399 | **0.0001** |
| I - T | -0.069206 | 0.005582 | 84 | -12.398 | **<.0001** |
| M - T | -0.083397 | 0.005499 | 84 | -15.166 | **<.0001** |
| Opsin = LWF2 |  |  |  |  |  |
| contrast | estimate | SE | df | t.ratio | p.value |
| I - M | -0.075228 | 0.035757 | 84 | -2.104 | 0.0951 |
| I - T | 0.378218 | 0.061868 | 84 | 6.113 | **<.0001** |
| M - T | 0.453446 | 0.060945 | 84 | 7.44 | **<.0001** |
| Opsin = LWF3 |  |  |  |  |  |
| contrast | estimate | SE | df | t.ratio | p.value |
| I - M | 0.003451 | 0.004212 | 84 | 0.819 | 0.6923 |
| I - T | -0.034753 | 0.007288 | 84 | -4.768 | **<.0001** |
| M - T | -0.038203 | 0.00718 | 84 | -5.321 | **<.0001** |
| Opsin = LWF4 |  |  |  |  |  |
| contrast | estimate | SE | df | t.ratio | p.value |
| I - M | 0.049729 | 0.023984 | 84 | 2.073 | 0.1016 |
| I - T | -0.212204 | 0.041498 | 84 | -5.114 | **<.0001** |
| M - T | -0.261933 | 0.040879 | 84 | -6.408 | **<.0001** |

**Table S9.** Just noticeable differences (JNDs) for three possible trichromatic visual systems based on electroretinogram (ERG) recordings from *I. heterosticta* (UV 360nm, SW 450nm, LW 525 nm)*, I. elegans* LWF1 opsin sensitivity (UV 360nm, SW 413 nm, LW 531 nm), and *I. elegans* LWF2 opsin sensitivity (UV 360nm, SW 413 nm, LW 543 nm) determined via heterologous expression in HEK293T cells. The SWS sensitivity for the LWF1 and LWF2 visual systems is the average lambda max value recorded from heterologous expression assays of both SWS opsins in the current study and the UVS lambda max is from the *I. heterosticta* ERG recordings in all models. Stimuli shows the visual stimuli being compared (A = androchrome female thorax reflectance, I = infuscans female thorax reflectance). Task indicates a possible function for each comparison. A JND value above 1 indicates that the probed visual system can distinguish both visual stimuli based on the model parameters. Differences between the LWF2 visual system, ERG, and LWF1 visual systems are shown in ΔJND_ERG_ and ΔJND_F1_, respectively.

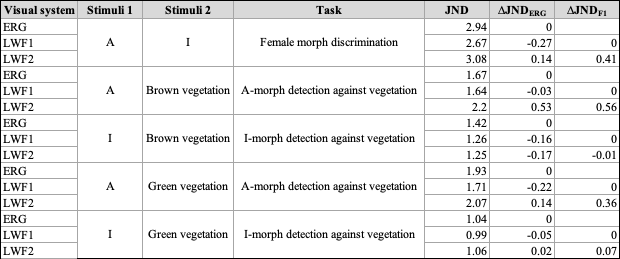

**Table S10.** Primer sequences used for amplification of *I. elegans* visual opsins, ocelli opsins, and housekeeping genes.

| **Oligo Name** | **Sequence (5’ tp 3’)** |
| --- | --- |
| Iel_UV_111F_1 | TGTGGGTGGGAAATATCTTGGG |
| Iel_UV_270R_1 | GATTCCATTCCCTGAAAGTGCG |
| Iel_SWb1_NR421F_1 | CGACATATACGGCTTGATTGGG |
| Iel_SWb1_571R_1 | ACCAAGTGAAGATGACGAGTCC |
| Iel_SWb2_786F_1 | GAAGATGCTCAAAGAACAGGCC |
| Iel_SWb2_1083R_1 | CACCTTCTTCTGGATCTCAGCC |
| Iel_LWA2_382F_1 | CTGCTACTTCGAGACATGGTCC |
| Iel_LWA2_595R_1 | AGTCCAGATGGCAGAGTTAAGC |
| Iel_LWD1_53F_1 | GTGGAAATTCCTTCTCCAACGC |
| Iel_LWD1_192R_1 | AGCCCACGAGATGAAACCG |
| Iel_LWE1_979F_2 | GTATCACAATGACTCCTCTCGTG |
| Iel_LWE1_1067R_2 | GACGCTTGAATCTGCTGCTC |
| Iel_LWF1_2F_1 | TGGAGACGGCAATGGTTCC |
| Iel_LWF1_200R_1 | TGACGAATGCCAAGATTCCATG |
| Iel_LWF2_908F_2 | CCATGACCATCAGCCCACT |
| Iel_LWF2_1110R_2 | CTGGCTGCTCGTCACCTC |
| Iel_LWF3_38F_2 | GGTCAACCCGAGGCTGC |
| Iel_LWF3_136R_2 | TCAAGTGCAACATTTCCGGA |
| Iel_LWF4_NR180F_2 | CGGTTGCTTGGGAATTGTGTC |
| Iel_LWF4_NR343R_2 | CCCAAATACCCAAGTTTCGTGATG |
| Iel_GADPH_5295F_1 | TATGATGCCTCCTTTTCCAGCC |
| Iel_GADPH_6025R_1 | GCCTTATGACCACTGTTCATGC |
| Iel_RL14_3641F_1 | ATCGTATCTGGCGTCTTTCTCG |
| Iel_RL14_4337R_1 | GCCACAACAAAGTCAGAGAACC |

**Table S11.** Post-hoc comparisons with Tukey correction showing differences in log relative opsin expression for teneral male *I. elegans* from two natural sites (Bunkeflostrand, Ilstorp) and the mesocosm tanks at Stensoffa Field Station. Significant differences are shown in bold. The final model included individual ID and replicate as random effects and showed a significant opsin x site interaction (df = 14, F = 8.5, *p* < 0.001) for teneral males.

| Opsin = SW1: | | | | | |
| --- | --- | --- | --- | --- | --- |
| contrast | estimate | SE | df | t.ratio | p.value |
| Bunkeflostrand - Ilstorp | 0.0256 | 0.142 | 9.81 | 0.18 | 0.9822 |
| Bunkeflostrand - Stensoffa | 0.1653 | 0.142 | 9.81 | 1.166 | 0.4986 |
| Ilstorp - Stensoffa | 0.1398 | 0.127 | 9.81 | 1.102 | 0.5346 |
| Opsin = SW2: | | | | | |
| contrast | estimate | SE | df | t.ratio | p.value |
| Bunkeflostrand - Ilstorp | -0.0451 | 0.142 | 9.81 | 0.318 | 0.946 |
| Bunkeflostrand - Stensoffa | 0.2927 | 0.142 | 9.81 | 2.065 | 0.1482 |
| Ilstorp - Stensoffa | 0.3378 | 0.127 | 9.81 | 2.664 | 0.0575 |
| Opsin = LWA: | | | | | |
| contrast | estimate | SE | df | t.ratio | p.value |
| Bunkeflostrand - Ilstorp | -0.0434 | 0.142 | 9.81 | 0.306 | 0.95 |
| Bunkeflostrand - Stensoffa | 0.5517 | 0.142 | 9.81 | 3.892 | **0.0079** |
| Ilstorp - Stensoffa | 0.5951 | 0.127 | 9.81 | 4.693 | **0.0023** |
| Opsin = LWF1: | | | | | |
| contrast | estimate | SE | df | t.ratio | p.value |
| Bunkeflostrand - Ilstorp | 0.0597 | 0.142 | 9.81 | 0.421 | 0.9077 |
| Bunkeflostrand - Stensoffa | 0.4521 | 0.142 | 9.81 | 3.189 | **0.0245** |
| Ilstorp - Stensoffa | 0.3924 | 0.127 | 9.81 | 3.095 | **0.0285** |
| Opsin = LWF2: | | | | | |
| contrast | estimate | SE | df | t.ratio | p.value |
| Bunkeflostrand - Ilstorp | 0.0038 | 0.142 | 9.81 | 0.027 | 0.9996 |
| Bunkeflostrand - Stensoffa | 0.7248 | 0.142 | 9.81 | 5.113 | **0.0013** |
| Ilstorp - Stensoffa | 0.721 | 0.127 | 9.81 | 5.686 | **0.0006** |
| Opsin = LWF3: | | | | | |
| contrast | estimate | SE | df | t.ratio | p.value |
| Bunkeflostrand - Ilstorp | 0.0679 | 0.142 | 9.81 | 0.479 | 0.8826 |
| Bunkeflostrand - Stensoffa | 0.5956 | 0.142 | 9.81 | 4.201 | **0.0049** |
| Ilstorp - Stensoffa | 0.5277 | 0.127 | 9.81 | 4.162 | **0.0052** |
| Opsin = LWF4: | | | | | |
| contrast | estimate | SE | df | t.ratio | p.value |
| Bunkeflostrand - Ilstorp | 0.0532 | 0.142 | 9.81 | 0.375 | 0.9258 |
| Bunkeflostrand - Stensoffa | 0.6435 | 0.142 | 9.81 | 4.539 | **0.0029** |
| Ilstorp - Stensoffa | 0.5903 | 0.127 | 9.81 | 4.655 | **0.0025** |
| Opsin = LWE: | | | | | |
| contrast | estimate | SE | df | t.ratio | p.value |
| Bunkeflostrand - Ilstorp | -0.1307 | 0.142 | 9.81 | 0.922 | 0.6397 |
| Bunkeflostrand - Stensoffa | -0.1412 | 0.142 | 9.81 | 0.996 | 0.596 |
| Ilstorp - Stensoffa | -0.0105 | 0.127 | 9.81 | 0.083 | 0.9962 |

**Table S12.** Post-hoc analysis with Tukey correction showing significant differences between relative opsin expression within teneral male *I. elegans*, excluding males from the Stensoffa Field station. Significant differences are shown in bold and comparisons that differ from analysis when including Stensoffa males are shown in italics (see main text and Table S4 for results including males from Stensoffa).

| Teneral | | | | | |
| --- | --- | --- | --- | --- | --- |
| contrast | estimate | SE | df | t.ratio | p.value |
| LWA - LWE | 1.529044 | 0.2123 | 1144 | 7.201 | **<.0001** |
| LWA - LWF1 | 0.23399 | 0.2123 | 1144 | 1.102 | 0.9565 |
| LWA - LWF2 | -0.088827 | 0.2123 | 1144 | -0.418 | 0.9999 |
| LWA - LWF3 | 0.233795 | 0.2123 | 1144 | 1.101 | 0.9567 |
| *LWA - LWF4* | *-0.61771* | *0.2123* | *1144* | *-2.909* | *0.0717* |
| LWA - SW1 | 0.994056 | 0.2123 | 1144 | 4.681 | **0.0001** |
| LWA - SW2 | 1.222805 | 0.2123 | 1144 | 5.759 | **<.0001** |
| LWE - LWF1 | -1.295054 | 0.2123 | 1144 | -6.099 | **<.0001** |
| LWE - LWF2 | -1.617871 | 0.2123 | 1144 | -7.619 | **<.0001** |
| LWE - LWF3 | -1.295249 | 0.2123 | 1144 | -6.1 | **<.0001** |
| LWE - LWF4 | -2.146754 | 0.2123 | 1144 | -10.11 | **<.0001** |
| LWE - SW1 | -0.534988 | 0.2123 | 1144 | -2.519 | 0.1882 |
| LWE - SW2 | -0.306239 | 0.2123 | 1144 | -1.442 | 0.8374 |
| LWF1 - LWF2 | -0.322817 | 0.2123 | 1144 | -1.52 | 0.7966 |
| LWF1 - LWF3 | -0.000195 | 0.2123 | 1144 | -0.001 | 1 |
| LWF1 - LWF4 | -0.8517 | 0.2123 | 1144 | -4.011 | **0.0017** |
| LWF1 - SW1 | 0.760065 | 0.2123 | 1144 | 3.579 | **0.0086** |
| LWF1 - SW2 | 0.988815 | 0.2123 | 1144 | 4.657 | **0.0001** |
| LWF2 - LWF3 | 0.322622 | 0.2123 | 1144 | 1.519 | 0.7971 |
| *LWF2 - LWF4* | *-0.528883* | *0.2123* | *1144* | *-2.491* | *0.2003* |
| LWF2 - SW1 | 1.082883 | 0.2123 | 1144 | 5.1 | **<.0001** |
| LWF2 - SW2 | 1.311632 | 0.2123 | 1144 | 6.177 | **<.0001** |
| LWF3 - LWF4 | -0.851505 | 0.2123 | 1144 | -4.01 | **0.0017** |
| LWF3 - SW1 | 0.760261 | 0.2123 | 1144 | 3.58 | **0.0085** |
| LWF3 - SW2 | 0.98901 | 0.2123 | 1144 | 4.657 | **0.0001** |
| LWF4 - SW1 | 1.611766 | 0.2123 | 1144 | 7.59 | **<.0001** |
| LWF4 - SW2 | 1.840515 | 0.2123 | 1144 | 8.667 | **<.0001** |
| SW1 - SW2 | 0.228749 | 0.2123 | 1144 | 1.077 | 0.9615 |

**Table S13.** Post-hoc analysis for relative opsin expression with Tukey correction between teneral (T), immature (I), and mature (M) male *I. elegans* for each opsin class, excluding males from Stensoffa Field station. Significant differences are shown in bold and comparisons that differ from analysis when including Stensoffa males are shown in italics (see main text and Table S7 for results including males from Stensoffa).

| Opsin = SW1: | | | | | |
| --- | --- | --- | --- | --- | --- |
| contrast | estimate | SE | df | t.ratio | p.value |
| I - M | -0.1156 | 0.0918 | 426 | -1.259 | 0.4191 |
| I - T | 0.2503 | 0.1956 | 414 | 1.279 | 0.4076 |
| *M - T* | *0.3659* | *0.1941* | *410* | *1.885* | *0.1443* |
| Opsin = SW2: | | | | | |
| contrast | estimate | SE | df | t.ratio | p.value |
| I - M | -0.0317 | 0.0918 | 426 | -0.345 | 0.9363 |
| I - T | 0.066 | 0.1956 | 414 | 0.338 | 0.9391 |
| M - T | 0.0978 | 0.1941 | 410 | 0.504 | 0.8696 |
| Opsin = LWA: | | | | | |
| contrast | estimate | SE | df | t.ratio | p.value |
| I - M | -0.0381 | 0.0918 | 426 | -0.415 | 0.9096 |
| I - T | -0.2956 | 0.1956 | 414 | -1.511 | 0.2866 |
| M - T | -0.2576 | 0.1941 | 410 | -1.327 | 0.381 |
| Opsin = LWF1: | | | | | |
| contrast | estimate | SE | df | t.ratio | p.value |
| I - M | 0.2238 | 0.0928 | 440 | 2.411 | **0.0431** |
| I - T | -0.9107 | 0.1957 | 414 | -4.653 | **<.0001** |
| M - T | -1.1345 | 0.1944 | 412 | -5.835 | **<.0001** |
| Opsin = LWF2: | | | | | |
| contrast | estimate | SE | df | t.ratio | p.value |
| I - M | -0.2718 | 0.0918 | 426 | -2.96 | **0.0091** |
| I - T | 0.5497 | 0.1956 | 414 | 2.81 | **0.0143** |
| M - T | 0.8215 | 0.1941 | 410 | 4.232 | **0.0001** |
| Opsin = LWF3: | | | | | |
| contrast | estimate | SE | df | t.ratio | p.value |
| I - M | -0.0686 | 0.0918 | 426 | -0.748 | 0.7353 |
| I - T | -0.3197 | 0.1956 | 414 | -1.634 | 0.2323 |
| M - T | -0.2511 | 0.1941 | 410 | -1.293 | 0.3997 |
| Opsin = LWF4: | | | | | |
| contrast | estimate | SE | df | t.ratio | p.value |
| I - M | -0.0695 | 0.0918 | 426 | -0.757 | 0.7297 |
| I - T | -0.2878 | 0.1956 | 414 | -1.471 | 0.3059 |
| M - T | -0.2183 | 0.1941 | 410 | -1.124 | 0.4995 |
| Opsin = LWE: | | | | | |
| contrast | estimate | SE | df | t.ratio | p.value |
| I - M | 0.0755 | 0.093 | 442 | 0.812 | 0.6957 |
| I - T | -0.5191 | 0.1956 | 414 | -2.654 | **0.0225** |
| M - T | -0.5946 | 0.1946 | 413 | -3.056 | **0.0067** |

**Table S14.** Post-hoc analysis with Tukey correction showing differences between proportion opsin expression between teneral (T), immature (I), and mature (M) male *I. elegans*, excluding males from the Stensoffa Field station. Significant differences are shown in bold and comparisons that differ from analysis when including Stensoffa males are shown in italics (see main text and Table S7 for results including males from Stensoffa).

| Opsin = SW1 | | | | | |
| --- | --- | --- | --- | --- | --- |
| contrast | estimate | SE | df | t.ratio | p.value |
| I - M | 1.88E-03 | 0.002421 | 81 | 0.778 | 0.7174 |
| I - T | 1.02E-02 | 0.005115 | 81 | 1.998 | 0.1193 |
| M - T | 8.33E-03 | 0.005064 | 81 | 1.646 | 0.2326 |
| Opsin = SW2 | | | | | |
| contrast | estimate | SE | df | t.ratio | p.value |
| I - M | 1.74E-03 | 0.000646 | 81 | 2.695 | **0.023** |
| I - T | 7.28E-05 | 0.001365 | 81 | 0.053 | 0.9984 |
| *M - T* | *-1.67E-03* | *0.001352* | *81* | *-1.235* | *0.4364* |
| Opsin = LWA | | | | | |
| contrast | estimate | SE | df | t.ratio | p.value |
| I - M | 4.23E-03 | 0.008493 | 81 | 0.498 | 0.8724 |
| I - T | -6.01E-02 | 0.017943 | 81 | -3.349 | **0.0035** |
| M - T | -6.43E-02 | 0.017764 | 81 | -3.621 | **0.0015** |
| Opsin = LWF1 | | | | | |
| contrast | estimate | SE | df | t.ratio | p.value |
| I - M | 1.42E-02 | 0.003121 | 81 | 4.547 | **0.0001** |
| I - T | -5.52E-02 | 0.006594 | 81 | -8.369 | **<.0001** |
| M - T | -6.94E-02 | 0.006528 | 81 | -10.627 | **<.0001** |
| Opsin = LWF2 | | | | | |
| contrast | estimate | SE | df | t.ratio | p.value |
| I - M | -7.52E-02 | 0.036373 | 81 | -2.068 | 0.1029 |
| I - T | 3.61E-01 | 0.076843 | 81 | 4.694 | **<.0001** |
| M - T | 4.36E-01 | 0.076077 | 81 | 5.73 | **<.0001** |
| Opsin = LWF3 | | | | | |
| contrast | estimate | SE | df | t.ratio | p.value |
| I - M | 3.45E-03 | 0.004275 | 81 | 0.807 | 0.6997 |
| I - T | -3.26E-02 | 0.009031 | 81 | -3.611 | **0.0015** |
| M - T | -3.61E-02 | 0.008941 | 81 | -4.033 | **0.0004** |
| Opsin = LWF4 | | | | | |
| contrast | estimate | SE | df | t.ratio | p.value |
| I - M | 4.97E-02 | 0.024393 | 81 | 2.039 | 0.1095 |
| I - T | -2.23E-01 | 0.051533 | 81 | -4.33 | **0.0001** |
| M - T | -2.73E-01 | 0.051019 | 81 | -5.348 | **<.0001** |

SUPPLEMENTAL FIGURES

**Figure S1.** Homology modelling of (A) *I. elegans* SWb1 opsin and (B) *I. elegans* LWF1 opsin obtained using the jumping spider crystal structure (PDB 6i9k) as template. The cis-retinal chromophore is colored in yellow. (A) Interacting residues predicted to lie within the binding pocket are colored in light grey, with two variants, candidate spectral residues, in blue (Y136, Y294, see also Table S3). (B) Residues predicted to interact with the chromophore are conserved between *I. elegans* LWF1 and LWF2, LWF3, LWF4 opsins, which is not unexpected given their close range of absorbance. Two variant residues between these 4 opsins and LWA2, are highlighted in yellow (A131, S319, see also Table S3). The templates and models were built in SwissModel followed by visualization in Pymol.

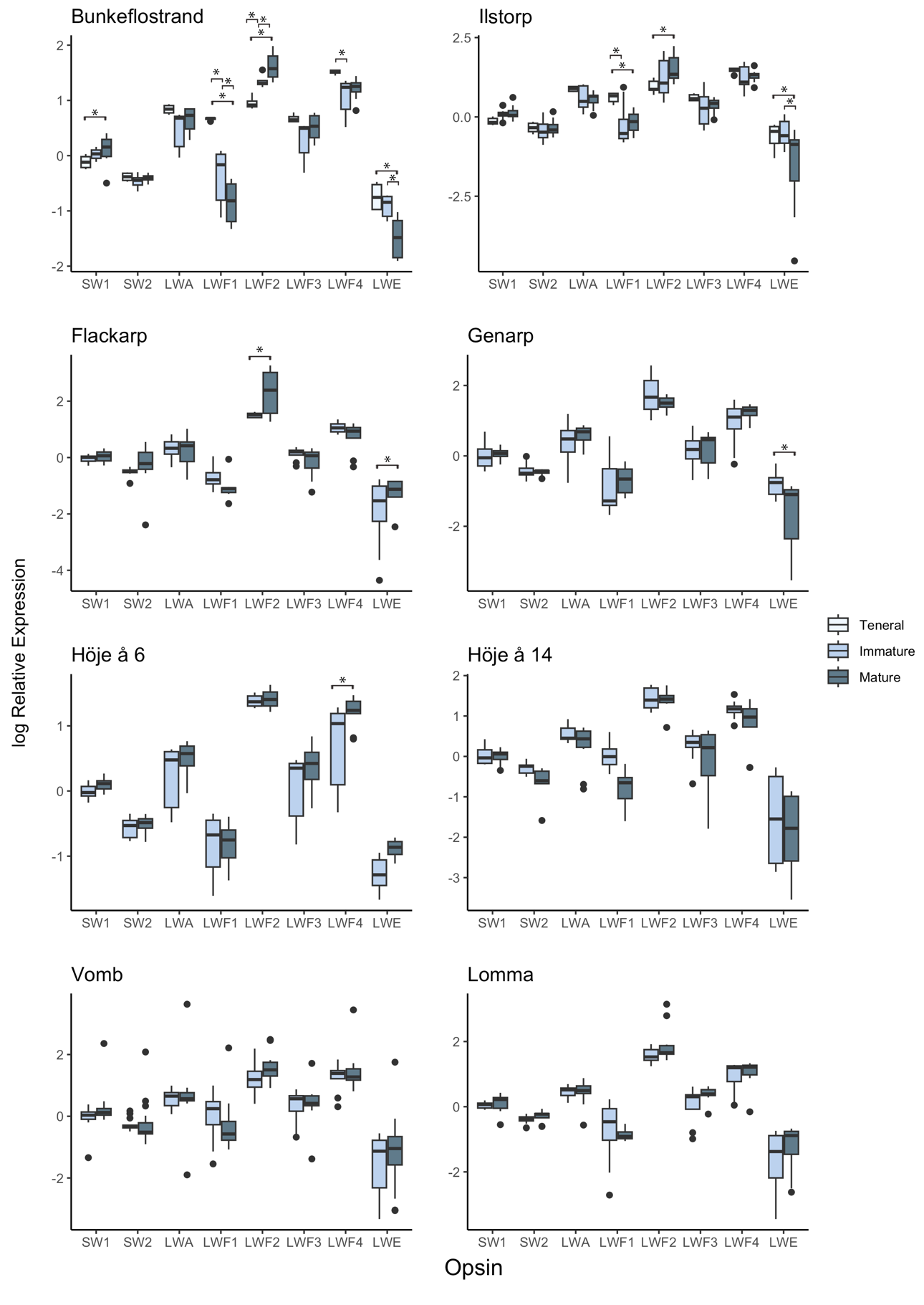

**Figure S2.** Opsin expression over adult development for male *I. elegans* from 8 sites across southern Sweden. Brackets indicate comparisons that are significant at an alpha level < 0.05. Bars in box and whisker plots show medians, boxes indicate upper and lower quartiles, whiskers show sample minima and maxima, and open circles show outliers. See Table S7 for p-values.

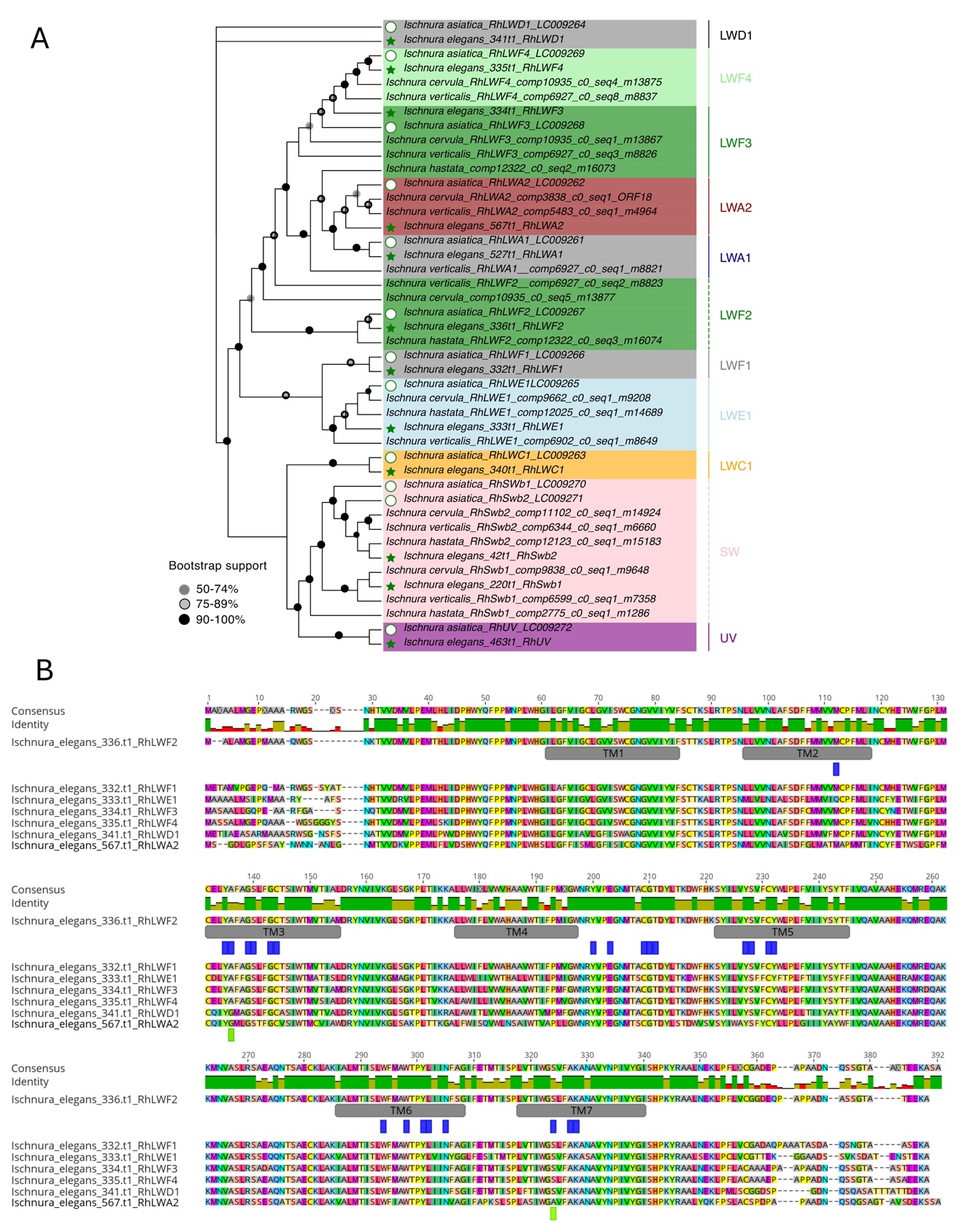
**Fig. S3. Opsin phylogeny and multiple alignment**. (A) *Ischnura* spp. opsin phylogeny displaying conserved opsin gene repertoires between *I. asiatica* (Futahashi et al., 2015) and *I. elegans* (sequences annotated in this study) (Price et al., 2022). Partial sequences for publicly available data of other *Ischnura* species were retrieved from NCBI. Depending on quality and completeness of available data, they may not represent the complete opsin paralogue set in those species. Opsin sequences were aligned in MAFFT (Katoh & Standley, 2013) and the phylogeny was reconstructed using IQtree (Nguyen et al., 2015) (Best-fit model LG+F+G4 with 1000 Ultrafast bootstrap replicates. The tree is unrooted with *Ischnura_asiatica_RhLWD1* drawn at root and was visualized with Evolview (Zhang et al., 2012). The *I. asiatica* and *I. elegans* orthologues for each opsin type, share between 91.5 % to 99.2% identity at the amino acid level, corresponding to 33 to 3 amino acid residue differences, *i.e.* *I. elegans-I. asiatica* pairwise comparisons (% identity amino acid level/number of substitutions)*:* UV: 91.5/33; SWb1: 96.1/15; SWb2: 93.4/26; LWA1:98.4/6; LWA2: 95/19; LWC1: 99.2/3; LWD1: 99.2/3; LWE1: 97.3/10; LWF1: 97.1/11; LWF2: 98.6/5; LWF3: 99.2/3; LWF4: 98.7/5). The multiple alignment and newick tree files are available as supplementary datasets accompanying the paper. (B) Multiple alignment of *I. elegans* LW opsins characterized in this study. Transmembrane domains are mapped with grey bars, based on homology modeling (Fig. S4), predicted residues lining the retinal binding pocket are indicated in blue, and two candidate spectral tuning sites between LWA2 and LWF opsins are indicated in light green (see also Table S4). The four LWF paralog opsins share 89-90% identity, and 63-69% with LWA2.

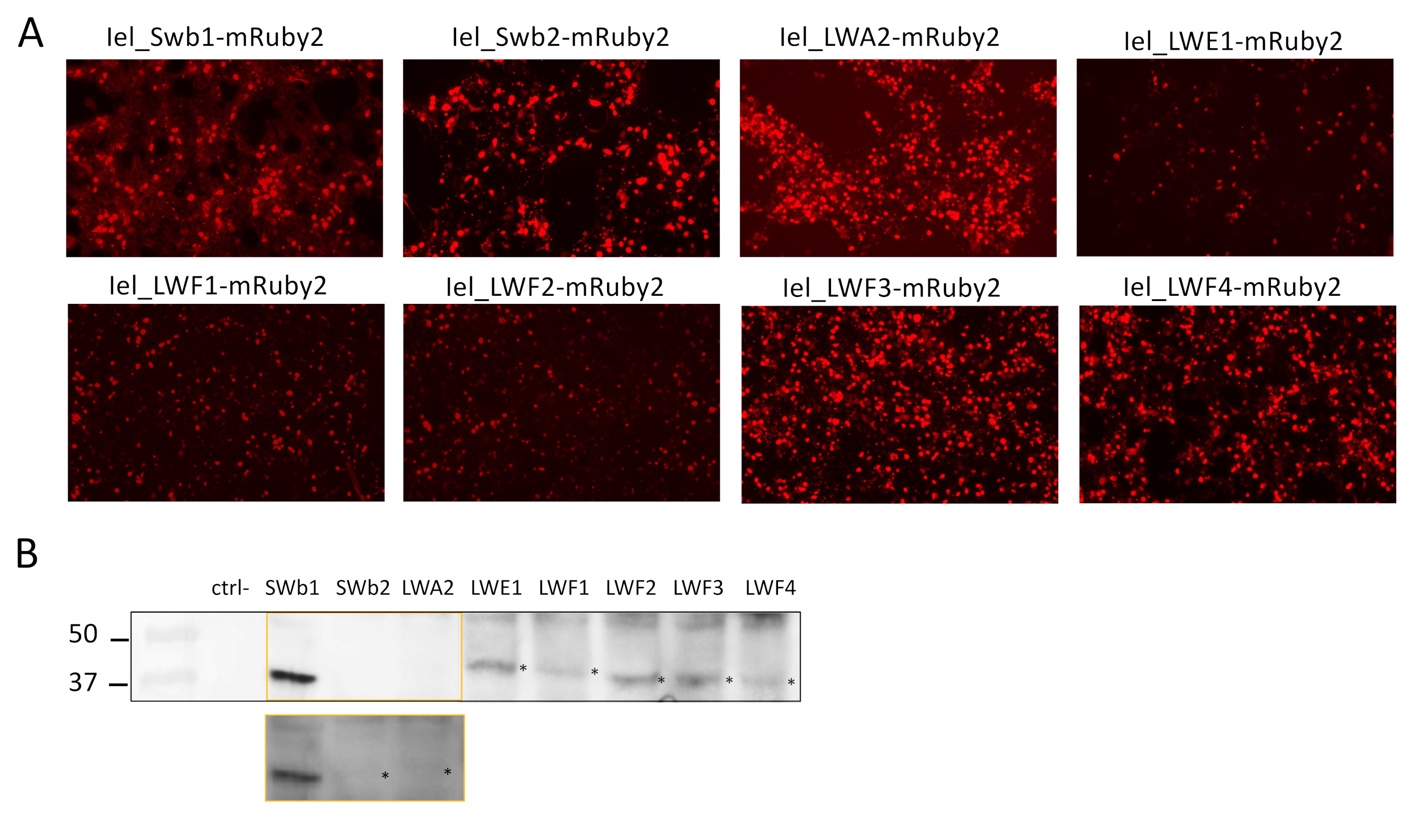

**Figure S4**. Small scale transfections verifying that each final pcDNA5-Opsin construct produced detectable protein levels in HEK293 cells. (A) Panels show transfected cells visualized under a fluorescent microscope to verify expression of mRuby2 indicating successful cloning of opsin open reading frames. (B) Western blot analysis and anti-FLAG signals of total protein lysates from a small scale 6-well culture dish assay, with lane 1: Kda ladder followed by untransfected HEK293T cells (ctrl-) followed by lysates from cell transfected with each opsin construct. Inset with increased contrast to detect lower protein expression of SWb2 and LWA2 opsins in the small-scale assay. Asterisks indicate predicted monomeric opsin units for each construct. Kda, ladder.

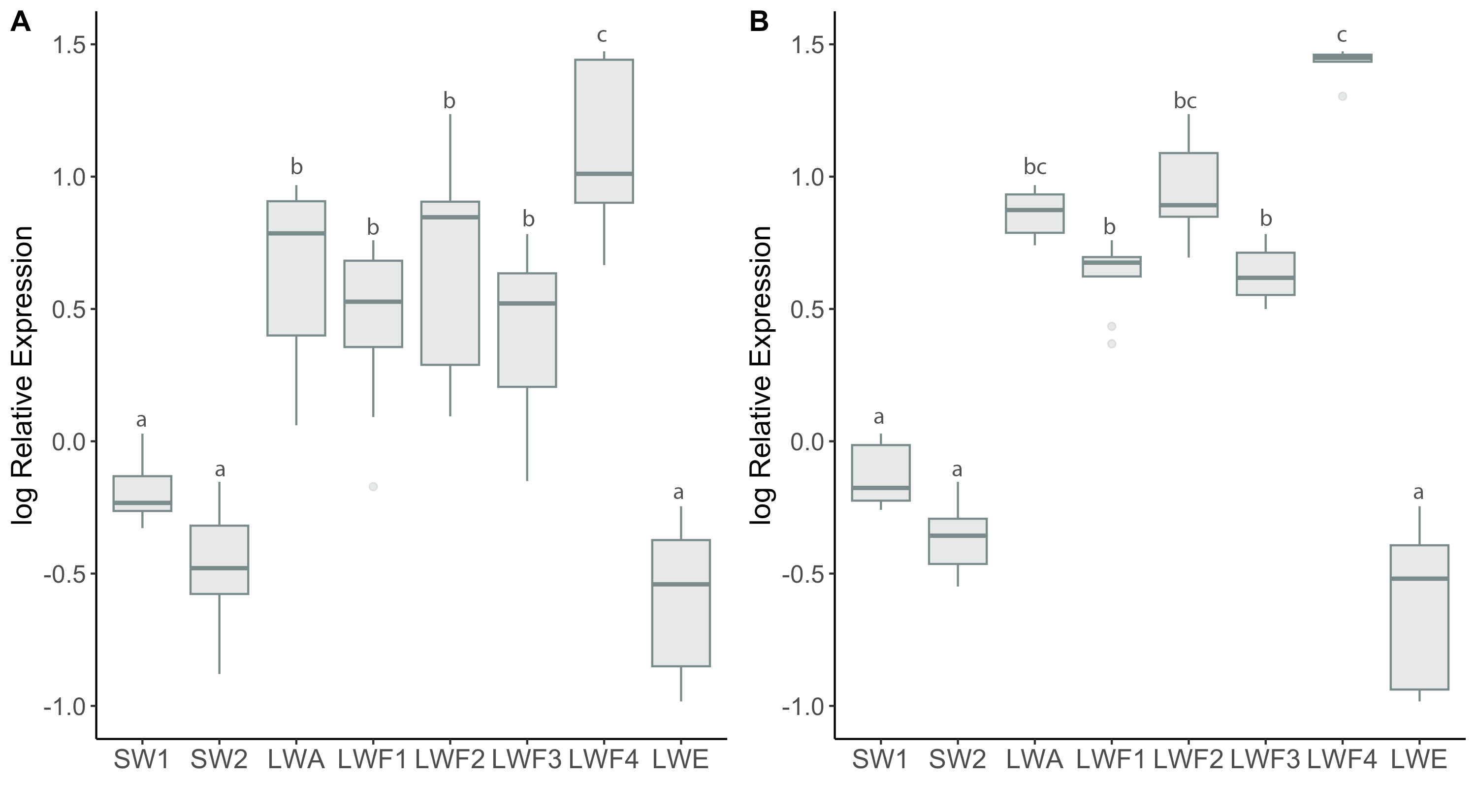

**Figure S5.** Relative opsin expression for (A) *I. elegans* teneral males from all three sampled locations and (B) teneral males from only natural sites (i.e., excluding Stensoffa)*.* Bars in box and whisker plots show medians, boxes indicate upper and lower quartiles, whiskers show sample minima and maxima, and open circles show outliers. Letters above each bar show significant differences between opsin types within each maturity level following Tukey corrections for multiple comparisons.
